# Engineering the bacterial cellulose-forming surface as a programmable protein recruitment interface

**DOI:** 10.64898/2026.07.13.738180

**Authors:** Shuang Zhang, Chengji Yang, Ruxia Fan, Sandra Kaabel, A. Sesilja Aranko, Markus B. Linder, Rahul Mangayil

**Affiliations:** Department of Bioproducts and Biosystems, School of Chemical Engineering, Aalto University, P.O. Box 16100, FI-02150 Espoo, Finland; Department of Chemistry and Material Sciences, School of Chemical Engineering, Aalto University, P.O. Box 16100, FI-02150 Espoo, Finland

## Abstract

Synthetic biology has advanced microorganisms to be programmed as production hosts, but its application to bacteria that inherently assemble extracellular materials remains limited. *Komagataeibacter* spp., natively synthesizes cellulose at the bacterial cell surface, creating a material-forming interface that has not been used as a programmable recruitment platform. Here we establish cell-surface display in *Komagataeibacter intermedius* and show that this interface can recruit defined proteins, making functionalization part of cellulose formation. By engineering Lpp’OmpA, we displayed a fluorescent protein and genetically encoded capture modules (SpyTag and SilkTag) to selectively capture catcher-fused protein cargos onto *K. intermedius* cell surface. Recruitment of silk-derived structural protein before cellulose production generated silk-associated fibrous structures within the pellicles, with retained cargo signal after washing. The resulting biocomposite showed reorganized fibre-network morphology, increased surface hydrophobicity, mesoscale ordering, and improved wet-state compressive strength. Wild-type cells exposed to same conditions did not reproduce these changes, demonstrating that material properties arise from surface-directed recruitment rather than protein exposure alone. This work demonstrates the material-forming bacterial surface as a programmable engineering interface for organizing extracellular proteins, providing a general strategy for engineering living materials.

## Introduction

Engineered living materials (ELMs) use genetically programmed cells to produce, organize or functionalize materials, offering a route to materials that can be grown under mild conditions and endowed with genetically encoded functions^1, 2^. *Komagataeibacter* spp. offer a distinct opportunity because they naturally synthesize bacterial cellulose (BC), with cellulose fibrils secreted and assembled near the cell surface^3, 4, 5^. This surface-proximal material formation represents an overlooked engineering opportunity, transforming *Komagataeibacter* from a conventional BC producer into a programmable chassis for biomolecule recruitment. Although genetic tools have been developed for *Komagataeibacter* spp., including modular cloning systems and engineered secretion strategies^6, 7^, the cellulose-producing cell surface remains underexplored as a programmable interface for spatially controlled protein recruitment during BC formation.

Existing BC functionalization strategies include post-synthetic adsorption, chemical modification and growth-associated functionalization^8^. Post-synthetic adsorption, and chemical modifications can introduce proteins or other functional components into pre-formed pellicles^9, 10^, albeit requiring additional downstream processing. ELMs approach instead functionalize BC during growth, either through engineered companion cells that secrete proteins into the forming matrix^11, 12, 13^, or by engineering the BC producer strain itself^7, 14^. However, secretion-based approaches require further process standardization and may limit scalability, whereas direct engineering of the BC producer has so far remained cargo-specific and may affect chassis fitness or BC synthesis^6, 15, 16, 17, 18^.

Cell-surface display offers an alternative route. Instead of individually expressing potential cargos in the BC producer, a capture module anchored on the living cell surface can recruit separately produced cargo at the site of fibril assembly, converting the bacterial cell surface into a programmable docking layer^19^. Among available surface-display scaffolds, Lpp’OmpA has been widely used to present heterologous proteins on Gram-negative bacterial surfaces^20, 21^. The Sec secretion and outer-membrane localization machinery required for Lpp’OmpA-mediated display is also present in *Komagataeibacter*^22, 23^, making this scaffold well suited for engineering the material-forming cell surface. However, direct fusion of each cargo to a surface anchor limits modularity, because every new cargo requires a separate construct, with varying expression, folding or display efficiencies^24, 25^. A more modular strategy is to display peptide modules facilitating the recruitment of individually expressed catcher-fused proteins^26^. SpyTag/SpyCatcher, a covalent peptide-protein coupling pair, has been used for protein immobilization on microbial cell surfaces^27, 28, 29^, while the recently developed SilkTag/SilkCatcher provides a recruitment pair designed for silk-derived structural proteins^26^. As displayed tags are short and genetically encoded, the same engineered surface can be combined with different catcher-fused cargos, separating surface programming from cargo selection^30^.

Here we engineer the material-forming cell surface of *Komagataeibacter intermedius* (*K. intermedius*) as a programmable protein recruitment interface. Using Lpp’OmpA as an outer-membrane anchoring platform, we displayed a directly fused fluorescent cargo (mLime) and used SpyTag to recruit SpyCatcher-mScarlet on *K. intermedius* cell surface. We then displayed SilkTag as a capture handle for a silk-derived structural cargo ^26, 31^.

Recruitment of this cargo to the cell surface before BC production yielded protein-associated pellicles with altered network morphology, mesoscale ordering and improved surface hydrophobicity and wet-state mechanics, changes not reproduced when wild-type (WT) cells were exposed to the same conditions. By programming recruitment at the site of nanocellulose assembly, this work demonstrates that material-forming bacterial surfaces can be repurposed to convert an inherent biomaterial producer into a chassis that organizes extracellular protein cargo into the material it forms.

## Results

### Establishing the cellulose-producing surface as a programmable recruitment interface

We engineered *K. intermedius* with an Lpp’OmpA display platform to present either directly fused protein cargos or short peptide capture modules for recruitment of catcher-fused proteins (Fig. 1A). To evaluate the platform, we generated constructs displaying (i) mLime protein, (ii) SpyTag peptide for covalent recruitment of SpyCatcher-fused proteins, and (iii) SilkTag peptide for the recruitment of a silk-derived structural cargo (SilkCatcher-rADF3-Sp-SpyTag). Each Lpp’OmpA fusion construct was expressed from a LuxR/P_lux_ inducible plasmid (Fig. 1B), and construct verification by agarose gel electrophoresis is shown in Fig. S1.

**Fig. 1.**
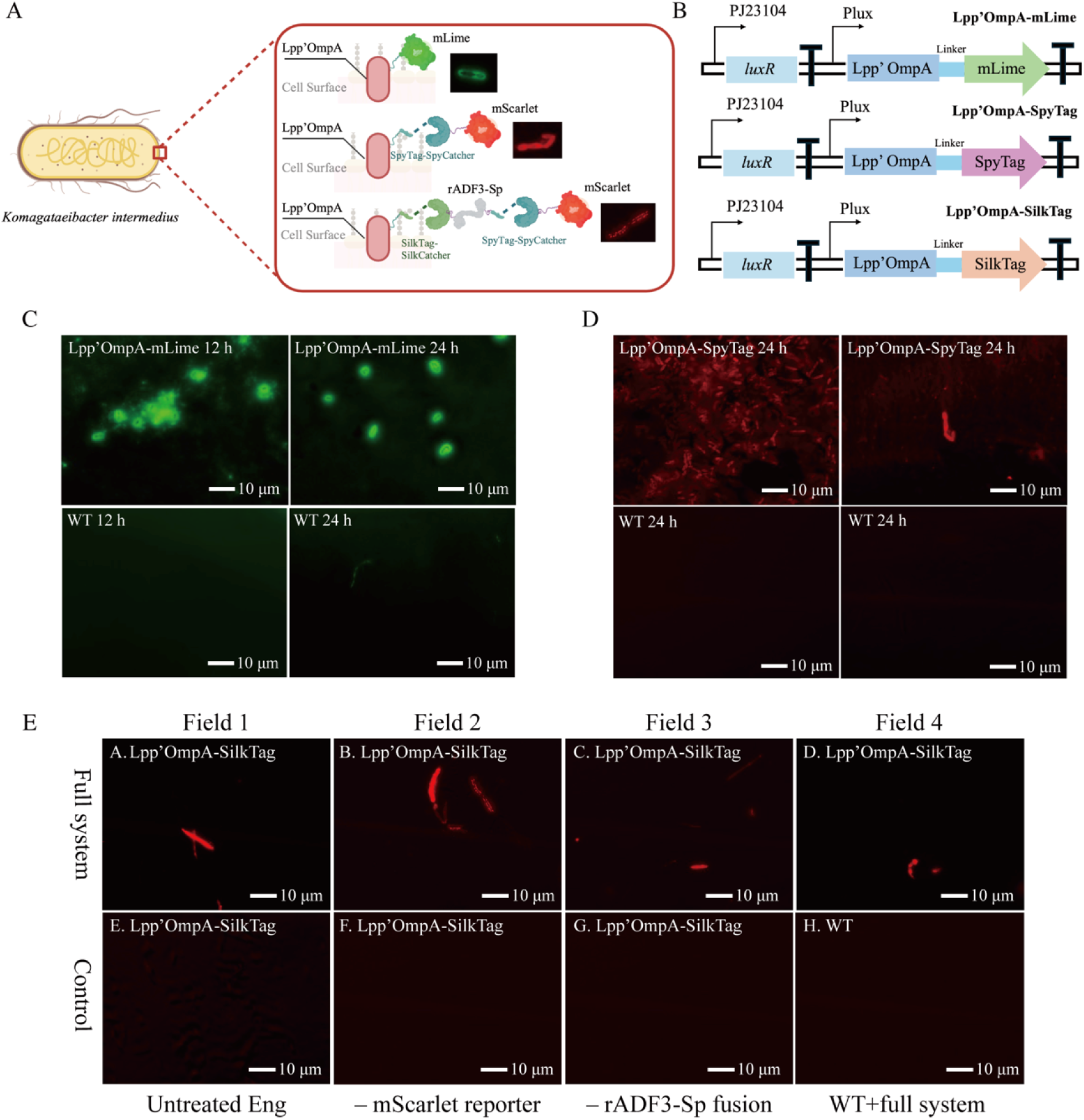
Establishment of a modular Lpp’OmpA-based surface-display platform in *K. intermedius*. **A**, Schematic of the Lpp’OmpA-based surface-display platform. Lpp’OmpA displays either a directly fused protein cargo (mLime) or peptide capture modules (SpyTag or SilkTag). SpyTag recruits SpyCatcher-mScarlet, whereas SilkTag recruits SilkCatcher-rADF3-Sp-SpyTag, which is subsequently labelled with SpyCatcher–mScarlet. **B**, Genetic architecture of the Lpp’OmpA display constructs, consisting of constitutive *luxR* expression and a P_lux_-controlled Lpp’OmpA fusion cassette encoding mLime, SpyTag or SilkTag. **C**, Representative fluorescence microscopy images of Lpp’OmpA-mLime cells and WT controls at 12 and 24 h. **D**, Representative fluorescence microscopy images showing SpyTag-mediated recruitment of SpyCatcher-mScarlet in Lpp’OmpA-SpyTag cells and WT controls. **E**, Recruitment of SilkCatcher–rADF3-Sp–SpyTag to Lpp’OmpA–SilkTag-expressing cells, visualized by subsequent labelling with SpyCatcher–mScarlet. Four representative fields are shown for the full system. Controls include untreated engineered cells, reactions lacking the mScarlet reporter or the rADF3-Sp fusion protein, and WT cells treated with the complete protein system. Images within each experiment were acquired under identical settings. Scale bars, 10 µm.

Cells expressing Lpp’OmpA-mLime exhibited a strong green, fluorescent signal, predominantly localized to the cell periphery, consistent with surface-associated localization, whereas WT cells displayed only background signal under the same imaging conditions (Fig. 1C). We next tested whether a displayed peptide tag remained accessible for protein recruitment. SpyTag-displaying cells retained SpyCatcher-mScarlet after washing, resulting in a strong red fluorescence signal, whereas WT cells treated with the same reporter exhibited minimal signal (Fig. 1D). Additional fluorescence microscopy images are provided in Figs. S2-3.

Finally, we tested whether this modular display strategy could recruit a functional structural protein cargo. We used rADF3-Sp, a spider-silk-derived protein with a propensity to self-assemble into β-sheet-rich materials^32^. SilkTag-displaying cells recruited SilkCatcher-rADF3-Sp-SpyTag, which was detected through terminal SpyTag labelling with SpyCatcher-mScarlet. Fluorescence signals were localized to the periphery of SilkTag-displaying cells, whereas untreated engineered cells, reactions lacking either the mScarlet reporter or rADF3-Sp fusion protein and WT cells treated with the complete protein system exhibited only background fluorescence (Fig. 1E).

Together, these results establish that the cellulose-producing surface is genetically addressable for modular protein recruitment.

### Surface-directed recruitment couples protein incorporation to cellulose formation

We next tested whether cell-surface recruitment could promote association of a silk-derived cargo with BC pellicles. SilkTag-displaying *K. intermedius* cells were incubated with SilkCatcher-rADF3-Sp-SpyTag prior to BC production, with WT cells exposed to same conditions as non-recruiting control (Fig. 2A). The resulting material groups are referred to throughout the study as WT-BC, WT-BC/rADF3-Sp, Eng-BC and Eng-BC/rADF3-Sp (Table S1).

**Fig. 2.**
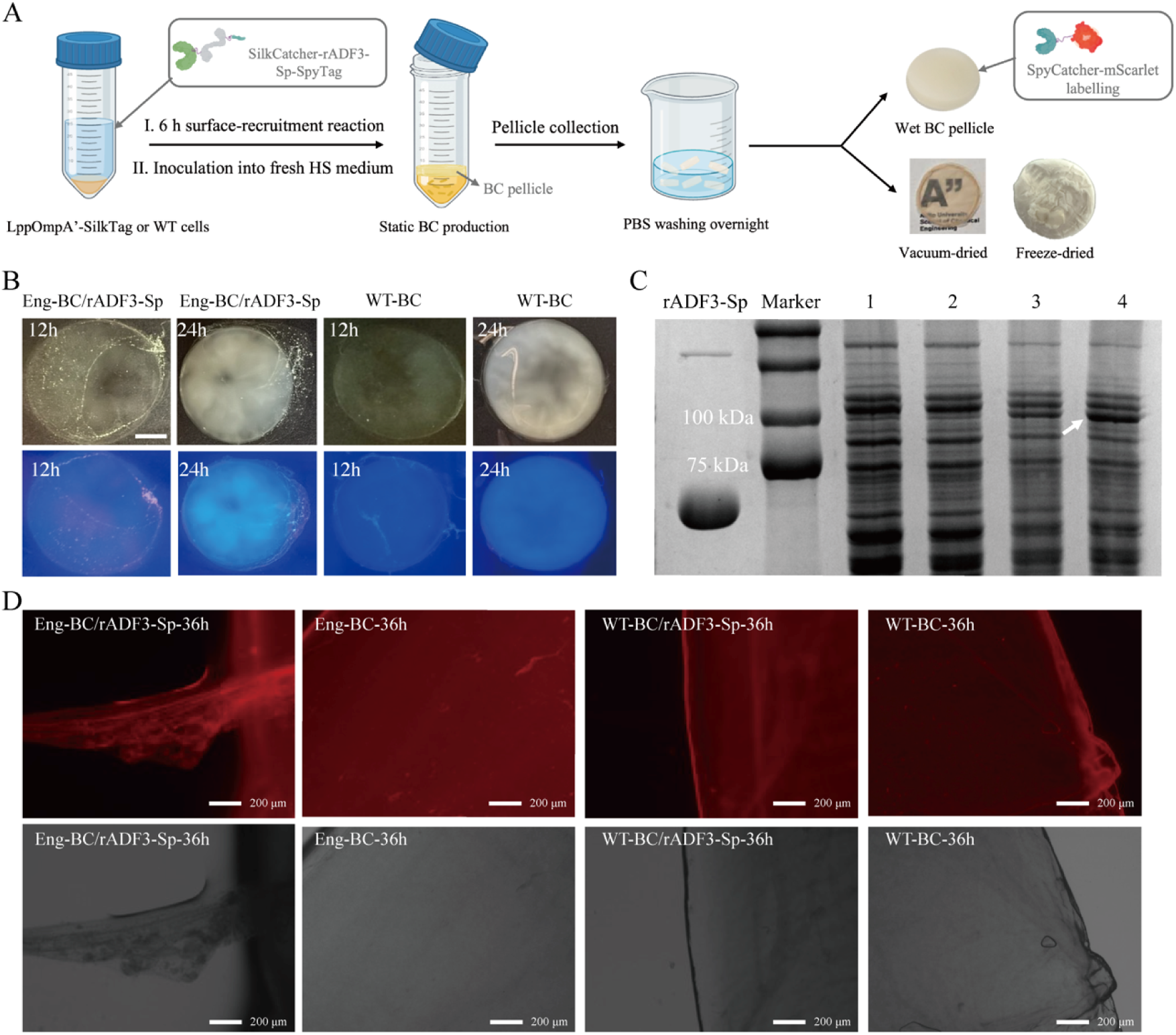
Formation of rADF3-Sp-associated BC pellicles through SilkTag-mediated recruitment. **A**, Schematic workflow for preparing and visualizing rADF3-Sp-associated bacterial cellulose pellicles. Lpp’OmpA-SilkTag-expressing or WT cells were incubated with SilkCatcher-rADF3-Sp-SpyTag for 6 h to allow surface recruitment and then inoculated into fresh HS medium for static bacterial cellulose production. The formed pellicles were collected, washed overnight in PBS and used as wet, vacuum-dried or freeze-dried materials for subsequent analysis. Wet pellicles were labelled with SpyCatcher-mScarlet for fluorescence imaging. **B**, Representative macroscopic images of bacterial cellulose pellicles formed at 12 and 24 h, shown under bright-field and fluorescence imaging conditions. **C**, SDS-PAGE analysis of pellicle-associated protein fractions. Purified SilkCatcher-rADF3-Sp-SpyTag was used as a reference. Lanes 1-4 correspond to WT-BC, WT-BC/rADF3-Sp, Eng-BC and Eng-BC/rADF3-Sp, respectively. The arrow indicates a protein band enriched in the Eng-BC/rADF3-Sp sample. **D,** Representative fluorescence microscopy images of washed 36 h pellicles after SpyCatcher-mScarlet labelling. Red fluorescence was more pronounced in Eng-BC/rADF3-Sp than in Eng-BC, WT-BC/rADF3-Sp and WT-BC controls. Scale bars, 200 µm.

Both engineered and WT cells produced BC pellicles, indicating that peptide display and rADF3-Sp recruitment did not impair pellicle formation (Fig. 2B). SDS–PAGE analysis of the pellicle-associated protein fraction revealed an enriched band in Eng-BC/rADF3-Sp that was absent from WT-BC/rADF3-Sp and the no-cargo controls (Fig. 2C). Notably, this band migrated at an apparent molecular weight higher than purified rADF3-Sp, consistent with the presence of a Lpp’OmpA-associated rADF3-Sp species rather than free rADF3-Sp alone. Consistent with these results, SpyCatcher-mScarlet labelling of washed pellicles showed stronger fluorescence signals in Eng-BC/rADF3-Sp than in Eng-BC, WT-BC/rADF3-Sp, or WT-BC (Fig. 2D). Additional images and full supporting data are provided in Figs. S4-S6.

Together, these results demonstrate that SilkTag-mediated recruitment promotes stable association of rADF3-Sp with BC pellicles produced by engineered *K. intermedius*.

### Surface-recruited rADF3-Sp imprints a protein-associated architecture and altered surface state into BC

Having established that SilkTag-mediated recruitment increased association of the silk-derived cargo in BC pellicles, we next examined whether this association altered pellicle organization chemical signatures and surface properties.

Because drying can strongly affect the appearance of hydrated BC networks^33, 34^, we compared the surface and fractured morphologies of Eng-BC/rADF3-Sp, WT-BC/rADF3-Sp, Eng-BC and WT-BC pellicles prepared by vacuum drying, air drying, freeze drying and liquid-nitrogen fracture using scanning electron microscopy (SEM) (Fig. 3A-E, Figs S7-10). Vacuum dried BC pellicles synthesized by Eng-BC/rADF3-Sp displayed pronounced aligned ridge-like surface features, whereas Eng-BC, WT-BC/rADF3-Sp and WT-BC showed irregular wrinkled morphology without comparable long-range organization (Fig. 3A, Fig. S7). A similar recruitment-dependent contrast was observed after ambient air drying (Fig. 3B, Fig. S8). In contrast, the freeze-dried Eng-BC/rADF3-Sp surface retained a more open fibrillar network with bridge-like features, suggesting that the aligned ridges observed after drying arise from drying-induced structural collapse (Fig. 3C, Fig. S9). Liquid-nitrogen-fractured Eng-BC/rADF3-Sp pellicles revealed locally connected fibrous features and bright bridge-like structures spanning adjacent cellulose-rich regions, whereas samples from WT-BC/rADF3-Sp, Eng-BC and WT-BC exhibited a more randomly entangled fibrillar network (Fig. 3D-E, Fig. S10). Although SEM morphology cannot determine the chemical composition of these structures, their selective appearance in Eng-BC/rADF3-Sp supports a distinct protein-associated architecture within the BC network.

**Fig. 3.**
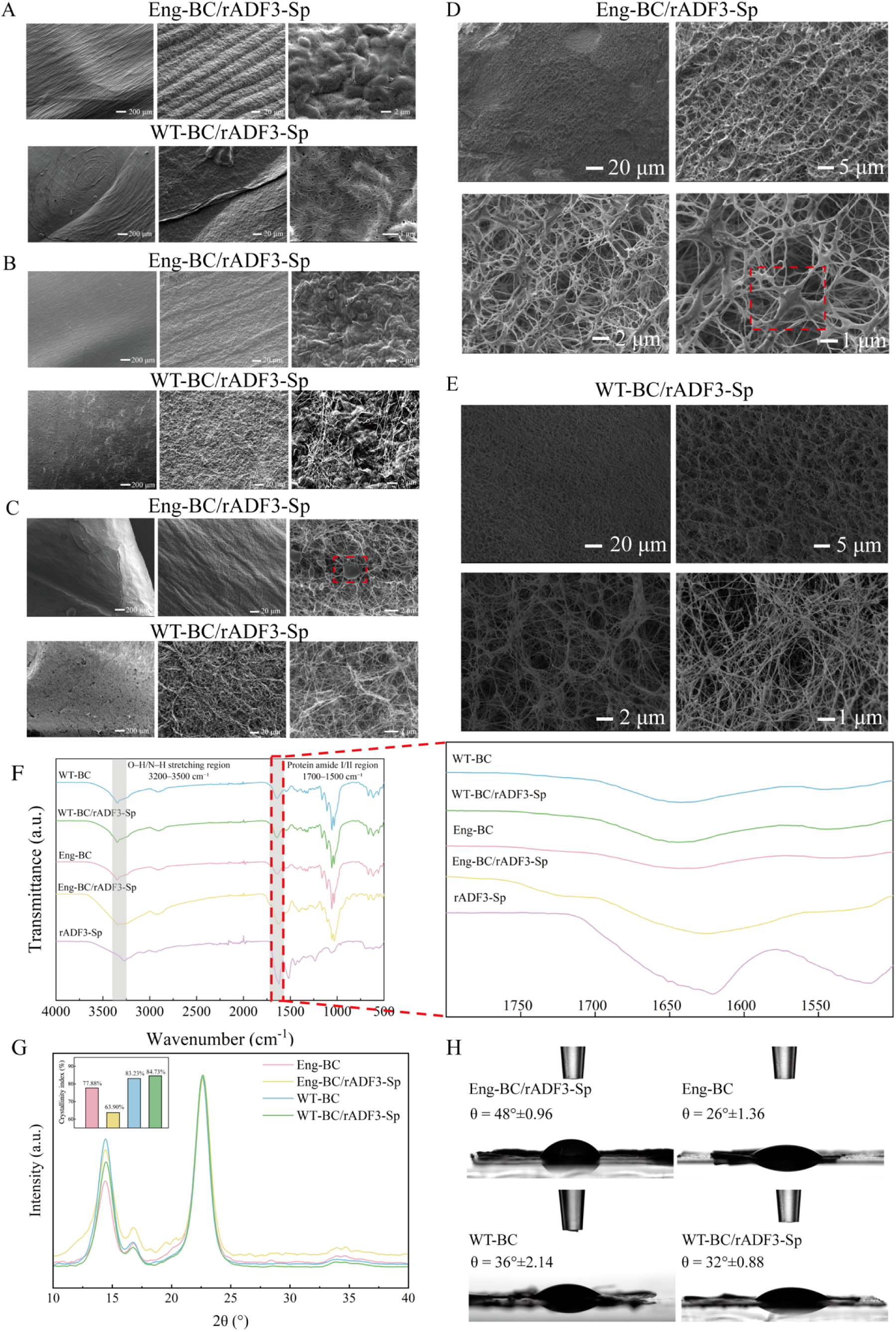
Surface-recruited rADF3-Sp generates a protein-associated architecture and altered surface state in bacterial cellulose pellicles. **A–C**, SEM images of Eng-BC/rADF3-Sp and WT-BC/rADF3-Sp pellicle surfaces prepared by vacuum drying (**A**), ambient air drying (**B**) and freeze drying (**C**). Eng-BC/rADF3-Sp showed aligned ridge-like surface structures after vacuum and air drying, whereas WT-BC/rADF3-Sp showed more irregular or randomly organized morphologies. The aligned ridge-like morphology was less evident after freeze drying. **D,E**, SEM images of liquid-nitrogen-fractured Eng-BC/rADF3-Sp (**D**) and WT-BC/rADF3-Sp (**E**) interfaces. Eng-BC/rADF3-Sp showed locally connected fibrous features and bright bridge-like structures within the fractured network, whereas these features were less evident in WT-BC/rADF3-Sp. Red dashed boxes highlight representative bridge-like features observed in Eng-BC/rADF3-Sp. **F**, FTIR spectra with an enlarged view of the amide-associated region. **G**, XRD patterns of the four BC pellicle groups; the inset shows the calculated apparent crystallinity index values. **H**, Representative water contact angle images of dried pellicles. Eng-BC/rADF3-Sp showed a higher contact angle than Eng-BC, whereas WT-BC/rADF3-Sp remained close to WT-BC. Scale bars are indicated in the SEM images.

We next assessed protein-associated chemical and structural signatures. Fourier-transform infrared (FTIR) spectra were dominated by cellulose-associated bands^35^ in all samples. However, compared to Eng-BC, Eng-BC/rADF3-Sp exhibited increased absorbance in the amide I/II region (1700 cm^-1^-1500-cm^-1^), consistent with the spectrum of purified rADF3-Sp (Fig. 3F). This observation supports the presence of silk-protein-associated chemical signatures in BC pellicles synthesized by Eng-BC/rADF3-Sp^36, 37^

X-ray diffraction (XRD) spectra showed characteristic BC diffraction profiles^38^ in all samples, indicating that cell-surface engineering and rADF3-Sp recruitment preserved the cellulose crystalline framework (Fig. 3G). Nevertheless, the apparent crystallinity index decreased from 77.85% in Eng-BC to 62.90% in Eng-BC/rADF3-Sp, whereas WT-BC and WT-BC/rADF3-Sp showed comparable values (83.21% and 84.78%, respectively).

Polarized optical microscopy (POM) further supported altered mesoscale organization. Eng-BC/rADF3-Sp showed more evident birefringent and band-like optical features than the control groups, whereas such patterns were not observed in BC pellicles synthesized by WT-BC/rADF3-Sp despite exposed to rADF3-Sp protein (Fig. S11).

Thermogravimetric analysis showed degradation profiles characteristic of BC and consistent with previous reports^39^ (Fig. S12). Eng-BC/rADF3-Sp exhibited a slightly higher residual weight than Eng-BC (30.0% and 26.1%, respectively), consistent with the differences in material composition following rADF3-Sp recruitment.

We then measured water contact angles to determine whether rADF3-Sp recruitment altered the surface wettability of vacuum-dried pellicles^40^. Representative contact angle images are shown in Fig. 3H, and the corresponding quantification is provided in Fig. S13. Water contact angles increased from 26 ± 1.36° for Eng-BC to 48 ± 0.96° for Eng-BC/rADF3-Sp (***P < 0.001). In contrast, WT-BC/rADF3-Sp showed no significant difference from WT-BC, with contact angles of 32 ± 0.88° and 36 ± 2.14°, respectively.

Together, these analyses demonstrate that SilkTag-mediated recruitment of rADF3-Sp reorganizes the developing BC network, producing recruitment-dependent changes in pellicle organization and surface properties while preserving the cellulose crystalline framework.

### Surface-recruited rADF3-Sp improves wet-state compressive response

The recruitment-dependent structural and surface changes observed in Eng-BC/rADF3-Sp suggested that the engineered pellicles might demonstrate altered mechanical behavior under hydrated conditions. As BC pellicles function as water-rich fibrous networks, we evaluated their wet-state load-bearing response by unconfined compression.

The compressive stress-strain curves differed among the four groups (Fig. 4A). Eng-BC/rADF3-Sp exhibited higher compressive stress than Eng-BC across the tested strain range, with the difference becoming more pronounced at higher strains. Quantification at 30%, 50% and 60% strain confirmed higher compressive stress in Eng-BC/rADF3-Sp than in Eng-BC (Fig. 4B-D). The apparent compressive modulus, calculated from the initial low-strain region, was also higher in Eng-BC/rADF3-Sp (Fig. 4E). In contrast, WT-BC/rADF3-Sp showed mechanical responses comparable to those of WT-BC, both in the stress-strain curves and quantified compression parameters.

**Fig. 4.**
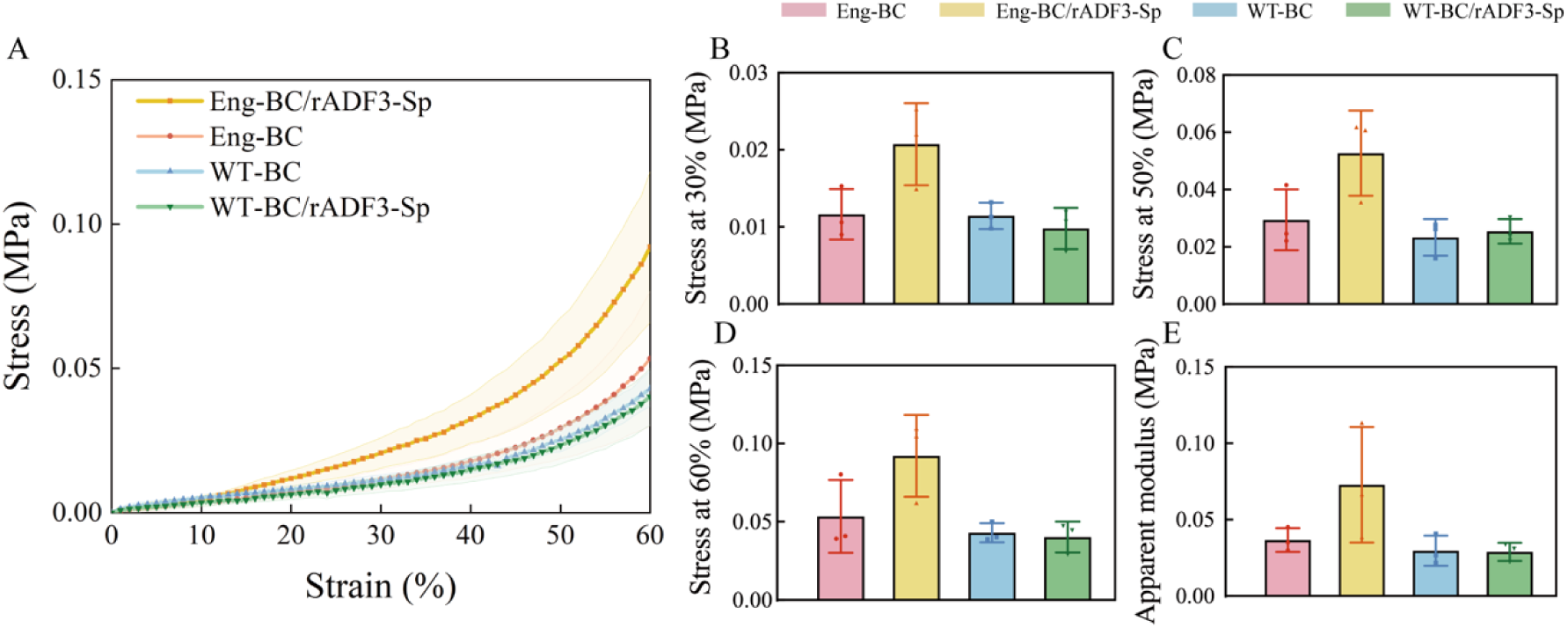
Wet-state compressive properties of rADF3-Sp-associated BC pellicles. **A**, Compressive stress-strain curves of hydrated BC pellicles under unconfined compression. Shaded regions indicate variation among independent samples. **B-D**, Compressive stress extracted from the stress-strain curves at 30%, 50% and 60% strain, respectively. **E**, Apparent compressive modulus calculated from the initial low-strain region of the stress-strain curves. Data are shown as mean ± SD, with individual points representing independent pellicles.

Together, these results demonstrate that surface-recruited rADF3-Sp improved the wet-state compressive properties of BC pellicles, consistent with the recruitment-dependent changes in pellicle architecture, composition and surface properties.

## Discussion

BC is typically valued as a high-purity extracellular polysaccharide, but its mode of biosynthesis also creates an underused engineering opportunity. Existing BC functionalization strategies generally rely on post-synthetic modification of purified pellicles or secretion of recombinant proteins into the culture medium, approaches that provide limited control over the spatial relationship between protein cargo and the developing cellulose network. In contrast, cellulose fibrils are secreted and assembled at the bacterial cell surface^41^, making *Komagataeibacter* spp. not only a biomaterial producing cell factory but also a spatial organizer of the nascent material. Here we used this feature to engineer the surface of *K. intermedius* as a genetically encoded interface for protein recruitment and BC functionalization.

The conceptual advance of this work is not surface display alone, or the addition of silk-derived protein to a *Komagataeibacter* culture. Rather, it is the repurposing of established surface display platform together with tag-catcher modules to link cell-surface engineering with extracellular material formation in an inherent biomaterial producer. The choice of Lpp’OmpA, a Sec-dependent display scaffold, leverages the conserved secretion machinery of *Komagataeibacter*, providing a simple and modular platform for programmable protein recruitment. Beyond demonstrating cell-surface presentation of proteins and peptides, this work shows that displayed capture modules can function as programmable handles that make the *Komagataeibacter* surface chemically addressable. This modular architecture decouples producer-cell engineering from cargo identity, allowing different catcher-fused proteins to be recruited without rebuilding the surface-display construct.

This modularity is particularly relevant for *Komagataeibacter* spp. because material assembly occurs close to the cell surface. In the rADF3-Sp experiment, the silk-derived fusion protein was recruited before BC production, positioning the cargo near the site of cellulose fibril secretion and assembly. The comparison with WT-BC/rADF3-Sp is therefore critical. Although both WT and engineered cultures were exposed to SilkCatcher-rADF3-Sp-SpyTag protein, only the engineered cells provided a defined SilkTag docking site. Consequently, the increased protein retention, together with the altered morphology, wettability and wet-state compressive response observed in Eng-BC/rADF3-Sp, indicate that the resulting material properties depended on how the cargo was presented at the cellulose-producing cell surface rather than on protein exposure alone. The apparent bridge-like features observed in liquid-nitrogen-fractured pellicles and the amide-associated FTIR signal further support the presence of a protein-associated architecture within the engineered BC network.

The mechanism by which surface-recruited rADF3-Sp alters BC remains to be resolved. We speculate that concentrating rADF3-Sp at the cellulose extrusion interface increases its local abundance during early network assembly, thereby modifying fibril–fibril interactions rather than cellulose biosynthesis itself. This interpretation is supported by the aligned surface features, altered fractured-interface morphology and reduced apparent crystallinity observed in pellicles produced by rADF3-Sp-recruiting cells. Additionally, the accompanying increase in water contact angle further indicates that these structural changes are reflected in the surface state of the resulting pellicle^42^, with Eng-BC/rADF3-Sp less wettable than Eng-BC and WT-BC/rADF3-Sp statistically similar to WT-BC. Collectively, these observations support a model in which localized protein recruitment modifies the organization of the developing cellulose network.

The wet-state compression data further link recruitment-dependent organizations to material performance. BC pellicles are hydrated fibrous networks, and their compressive response depends on fibre packing, interfibrillar interactions and water distribution, as well as on crystallinity^43, 44^. Eng-BC/rADF3-Sp showed higher compressive stress and apparent compressive modulus than Eng-BC, while WT-BC/rADF3-Sp remained close to WT-BC. Together, these findings suggest that surface-guided recruitment affected the load-bearing organization of the pellicle.

This study has several limitations that should be addressed in future work. The present study uses rADF3-Sp as a model structural cargo, and broader generality will require testing additional catcher-fused proteins with different sizes, functions and assembly behaviors. More direct measurements of cargo distribution during BC formation are also needed. Live imaging, immunolabelling, higher-resolution microscopy or quantitative mass spectrometry could help determine protein recruitment load, retention and distribution during pellicle growth and processing. These approaches should provide a more direct understanding of cargo recruitment, retention and distribution during pellicle formation.

In the main experiment, purified SilkCatcher-rADF3-Sp-SpyTag was used to establish the recruitment principle under controlled conditions. The preliminary lysate-compatibility experiment suggests a possible route toward more integrated production workflows (Fig. S14). Although this does not demonstrate direct recruitment from crude lysate, it suggests that *K. intermedius* can tolerate lysate-containing environments, which may facilitate future process development for scalable material fabrication^44^.

More broadly, this study reframes BC-producing bacteria as programmable organizers of extracellular materials. Although rADF3-Sp was used here as a model silk-derived structural cargo, the same recruitment logic could be applicable to other catcher-fused proteins, provided they remain functional after cell-surface localization. The modular nature of the platform should also facilitate the incorporation of diverse recombinant protein cargos without redesigning the engineered producer strain. Additionally, future work will determine whether simultaneous display of multiple orthogonal tag-catcher pairs enables spatially controlled recruitment of multiple protein functions within the same cellulose matrix. Rather than modifying BC after pellicle formation or utilizing model microbes to secrete functional proteins, this approach enables the material-forming cell surface itself to be engineered as a modular input point for protein function. These findings establish the material-forming bacterial surface as a programmable engineering interface for organizing extracellular biomolecules, providing a general strategy for constructing living materials with genetically encoded functions.

## Methods

### Strains and cultivation

*Escherichia coli* TOP10 (Thermo Fisher Scientific) was used for cloning and plasmid propagation. Recombinant *E. coli* strains were cultured in Luria-Bertani medium at 37 °C with appropriate antibiotics. Unless otherwise stated, *K. intermedius*^45^ was cultivated in Hestrin-Schramm medium (HS medium) containing 5 g L ¹ peptone, 5 g L ¹ yeast extract, 2.7 g L ¹ Na HPO, 1.5 g L ¹ citric acid and 2% (w/v) glucose at 30 °C. Unless otherwise stated, medium components and antibiotics were purchased from Sigma-Aldrich. Chloramphenicol and spectinomycin were used at final concentrations of 340 µg mL ¹ and 250 µg mL ¹, respectively.

### Plasmid construction and surface-display expression

Surface-display constructs were generated by fusing the Lpp’OmpA anchor to mLime, SpyTag or SilkTag under LuxR/P_lux_ inducible system. Plasmids were assembled by Golden Gate cloning^46^, propagated in *E. coli* TOP10 and verified by colony PCR and sequencing. Verified plasmids were introduced into *K. intermedius* by electroporation. For surface-display expression, recombinant *K. intermedius* cultures were grown in HS medium with 1 µg mL ¹ AHL^7^ (N-(β-ketocaproyl)-L-homoserine lactone; Sigma-Aldrich), collected by centrifugation, washed at least three times with sterile 1× PBS and resuspended in sterile 1× PBS for fluorescence microscopy or protein-binding assays. Genetic parts and primers used in this study are listed in Data S1 and Table S2.

### Protein-binding assays

For SpyTag validation, *K. intermedius* cells expressing surface-displayed SpyTag were incubated with purified SpyCatcher-mScarlet at a final concentration of 20.4 µM at 4 °C for 4 h. Cells were washed thrice with sterile 1× PBS before fluorescence microscopy^47^. For SilkTag-mediated recruitment, SilkTag displaying *K. intermedius* cells were incubated with purified SilkCatcher-rADF3-Sp-SpyTag, followed by SpyCatcher-mScarlet labelling. Reactions were performed in sterile 1× PBS at room temperature (RT) for 6 h, followed by repeated PBS washing before imaging. WT and untreated engineered cells and reactions lacking individual protein components were used as controls.

### Preparation of rADF3-Sp-associated BC pellicles

BC pellicles used for inoculum preparation were digested with 2% (w/v) cellulase (Sigma-Aldrich) to release the cells entrapped within the BC pellicle. Digested samples were washed with sterile 1× PBS, collected by centrifugation and weighed in the wet state.

### Pellicle washing and sample preparation

After static cultivation, pellicles were collected with sterile forceps and washed three times with sterile water or 1× PBS. For extensive washing, samples were incubated in PBS overnight. Washed pellicles were analyzed in the hydrated state or processed by air-drying, vacuum-drying at 50 °C or freeze-drying. For fractured-interface SEM imaging, dried samples were fractured after immersion in liquid nitrogen.

### Microscopy and material characterization

Fluorescence microscopy was performed using a Zeiss Axio Vert A1 inverted microscope with an mScarlet-compatible channel. SEM imaging was performed using a JEOL JSM-IT800HL after Au/Pd sputter 5 nm coating. Protein retention in washed pellicles was analyzed by SDS-PAGE after cellulase digestion, using purified SilkCatcher-rADF3-Sp-SpyTag as a reference. FTIR, XRD, TGA and polarized optical microscopy were performed as described in the Supplementary Methods.

### Mechanical testing

Hydrated pellicles were tested by unconfined compression using an Instron 4204 universal testing machine. Samples with an approximate height of 3 mm were compressed at 0.02 mm s ¹ after applying a 0.2 N preload. Unless otherwise stated, samples were compressed to 60% strain. The apparent compressive modulus was calculated from the 0-10% strain region.

### Water contact angle

Static water contact angles of vacuum-dried pellicles were measured using a Theta Flex optical tensiometer (Biolin Scientific). A water droplet was placed on the sample surface, and contact angles were calculated immediately after deposition. Three independent pellicles were measured per group.

### Statistical analysis

Data are presented as mean ± SD unless otherwise stated. Statistical analyses were performed using GraphPad Prism and Origin. Multiple-group comparisons were performed using one-way ANOVA followed by Tukey’s multiple-comparison test^48^ when assumptions of normality and equal variance were met. Unless otherwise stated, the numbers of biological and technical replicates are indicated in the corresponding Fig. legends.

## Supporting information

Supplementary Information for the manuscript

## Acknowledgements

This work was a part of the Ministry of Education and Culture’s Doctoral Education Pilot under Decision No. VN/3137/2024-OKM-6 (Circular Materials Bioeconomy Network, CIMANET). R.M. acknowledges the Research Council of Finland (Grant Number 346983) for supporting this study. The authors thank Ke Yao-jin for assistance with polarized optical microscopy measurements.

## Author information

Department of Bioproducts and Biosystems, School of Chemical Engineering, Aalto University, P.O. Box 16100, FI-02150 Espoo, Finland

Shuang Zhang, Ruxia Fan, A. Sesilja Aranko, Markus B. Linder & Rahul Mangayil

Department of Chemistry and Materials Science, School of Chemical Engineering, Aalto University, P.O. Box 16100, FI-02150 Espoo, Finland

Chengji Yang & Sandra Kaabel

## Author contributions

S.Z. contributed to experimental design, biological experiments and data organization. C.Y. contributed to material-related experiments and data collection. R.F. contributed to protein purification and manuscript revision. K.S., A.S.A., M.B.L. and R.M. supervised the study, provided conceptual guidance, contributed resources and reviewed the manuscript. S.Z. and C.Y wrote the manuscript with input from all authors. All authors reviewed and approved the final manuscript.

## Competing interests

The authors declare no competing interests.

## Data availability

The data supporting the findings of this study are available within the article and its Supplementary Information. Additional raw data are available from the corresponding author upon reasonable request.

## Materials availability

Plasmids, engineered bacterial strains and other materials generated in this study are available from the corresponding author upon reasonable request, subject to institutional and biosafety regulations.

## Ethics and biosafety statement

This study did not involve human participants or animal experiments. All work with genetically modified microorganisms was conducted in accordance with institutional biosafety guidelines.

## Supplementary information

Supplementary Information is available for this paper.

## Notes

### Competing Interest Statement

The authors have declared no competing interest.

## References

1. Wang, Y., et al. Engineered living materials (ELMs) design: From function allocation to dynamic behavior modulation. Current Opinion in Chemical Biology 70, 102188 10.1016/j.cbpa.2022.102188 (2022).

2. Nguyen, P. Q., Courchesne, N.-M. D., Duraj-Thatte, A., Praveschotinunt, P. & Joshi, N. S. Engineered Living Materials: Prospects and Challenges for Using Biological Systems to Direct the Assembly of Smart Materials. Advanced Materials 30, 1704847 10.1002/adma.201704847 (2018).

3. Bersanetti, D. & Mangayil, R. Exploring Genome-Wide Mutation Dynamics and Bacterial Cellulose Impairment in Komagataeibacter intermedius Cultivated Under Agitation Stress. Computational and Structural Biotechnology Journal 35, 0009 10.34133/csbj.0009.

4. Girard, V.-D., Chaussé, J. & Vermette, P. Bacterial cellulose: A comprehensive review. Journal of Applied Polymer Science 141, e55163 10.1002/app.55163 (2024).

5. Nicolas, W. J., Ghosal, D., Tocheva, E. I., Meyerowitz, E. M. & Jensen, G. J. Structure of the Bacterial Cellulose Ribbon and Its Assembly-Guiding Cytoskeleton by Electron Cryotomography. J Bacteriol 203, 10.1128/jb.00371-20 (2021).

6. Florea, M., et al. Engineering control of bacterial cellulose production using a genetic toolkit and a new cellulose-producing strain. Proceedings of the National Academy of Sciences 113, E3431–E3440 10.1073/pnas.1522985113 (2016).

7. Goosens, V. J., et al. Komagataeibacter Tool Kit (KTK): A Modular Cloning System for Multigene Constructs and Programmed Protein Secretion from Cellulose Producing Bacteria. ACS Synthetic Biology 10, 3422–3434 10.1021/acssynbio.1c00358 (2021).

8. Malcý, K., Li, I. S., Kisseroudis, N. & Ellis, T. Modulating Microbial Materials - Engineering Bacterial Cellulose with Synthetic Biology. ACS Synthetic Biology 13, 3857–3875 10.1021/acssynbio.4c00615 (2024).

9. Shishparenok, A. N., Furman, V. V., Dobryakova, N. V. & Zhdanov, D. D. Protein Immobilization on Bacterial Cellulose for Biomedical Application. Polymers (Basel) 16, 10.3390/polym16172468 (2024).

10. Chen, S., et al. Bio-orthogonal functionalization of bacterial cellulose combining metabolic glycoengineering and click chemistry. Nat Commun 17, 10.1038/s41467-026-69130-8 (2026).

11. Gilbert, C., et al. Living materials with programmable functionalities grown from engineered microbial co-cultures. Nature Materials 20, 691–700 10.1038/s41563-020-00857-5 (2021).

12. Paronyan, M., et al. Co-Cultivation of Komagataeibacter xylinus MS2530 with Various Yeast Strains: Production and characterization of bacterial cellulose Films. Carbohydrate Polymer Technologies and Applications 11, 100840 10.1016/j.carpta.2025.100840 (2025).

13. Brugnoli, M., et al. Co-cultivation of Komagataeibacter sp. and Lacticaseibacillus sp. strains to produce bacterial nanocellulose-hyaluronic acid nanocomposite membranes for skin wound healing applications. International Journal of Biological Macromolecules 299, 140208 10.1016/j.ijbiomac.2025.140208 (2025).

14. Vannas, J., et al. A signal peptide-guided platform for in situ functionalization of bacterial nanocellulose in Komagataeibacter rhaeticus. bioRxiv, 2025.2012.2015.694344 10.64898/2025.12.15.694344 (2026).

15. Cabo, M., Jr., et al. Bacterial Nanocellulose Functionalization for Smart Bioelectronics: Integration into Biosensing, Neural Interfaces, and Tissue Engineering. ACS Polymers Au 5, 723–755 10.1021/acspolymersau.5c00074 (2025).

16. Gilbert, C. & Ellis, T. Biological Engineered Living Materials: Growing Functional Materials with Genetically Programmable Properties. ACS Synthetic Biology 8, 1–15 10.1021/acssynbio.8b00423 (2019).

17. Lu, Y., Mehling, M., Huan, S., Bai, L. & Rojas, O. J. Biofabrication with microbial cellulose: from bioadaptive designs to living materials. Chemical Society Reviews 53, 7363–7391 10.1039/d3cs00641g (2024).

18. Wang, S., Zhan, Y., Jiang, X. & Lai, Y. Engineering Microbial Consortia as Living Materials: Advances and Prospectives. ACS Synthetic Biology 13, 2653–2666 10.1021/acssynbio.4c00313 (2024).

19. Löfblom, J. Bacterial display in combinatorial protein engineering. Biotechnol J 6, 1115–1129 10.1002/biot.201100129 (2011).

20. Earhart, C. F. Use of an Lpp-OmpA fusion vehicle for bacterial surface display. Methods Enzymol 326, 506–516 10.1016/s0076-6879(00)26072-2 (2000).

21. Yang, C., et al. Cell Surface Display of Functional Macromolecule Fusions on Escherichia coli for Development of an Autofluorescent Whole-Cell Biocatalyst. Environmental Science & Technology 42, 6105–6110 10.1021/es800441t (2008).

22. Rutherford, N. & Mourez, M. Surface display of proteins by Gram-negative bacterial autotransporters. Microbial Cell Factories 5, 22 10.1186/1475-2859-5-22 (2006).

23. Francisco, J. A., Earhart, C. F. & Georgiou, G. Transport and anchoring of beta-lactamase to the external surface of Escherichia coli. Proceedings of the National Academy of Sciences 89, 2713–2717 10.1073/pnas.89.7.2713 (1992).

24. Gallus, S., Mittmann, E. & Rabe, K. S. A Modular System for the Rapid Comparison of Different Membrane Anchors for Surface Display on Escherichia coli. ChemBioChem 23, e202100472 10.1002/cbic.202100472 (2022).

25. Lee, S. Y., Choi, J. H. & Xu, Z. Microbial cell-surface display. Trends in Biotechnology 21, 45–52 10.1016/S0167-7799(02)00006-9 (2003).

26. Fan, R., et al. Biomolecular Click Reactions Using a Minimal pH-Activated Catcher/Tag Pair for Producing Native-Sized Spider-Silk Proteins. Angewandte Chemie International Edition 62, e202216371 10.1002/anie.202216371 (2023).

27. Hatlem, D., Trunk, T., Linke, D. & Leo, J. C. Catching a SPY: Using the SpyCatcher-SpyTag and Related Systems for Labeling and Localizing Bacterial Proteins. Int J Mol Sci 20, 10.3390/ijms20092129 (2019).

28. Che, S., Konno, H. & Makabe, K. SpyTag Peptide with Alkoxyl Aspartic Acids for pH-Dependent Activation of the SpyCatcher/Tag System. Bioconjugate Chemistry 35, 616–622 10.1021/acs.bioconjchem.4c00052 (2024).

29. Cai, M.-m., et al. The SpyTag/SpyCatcher System: Precise Regulation of Covalent Conjugation and Expansion of Application Scenarios. Biotechnology Journal 20, e70131 10.1002/biot.70131 (2025).

30. Fan, R. & Aranko, A. S. Catcher/Tag Toolbox: Biomolecular Click-Reactions For Protein Engineering Beyond Genetics. ChemBioChem 25, e202300600 10.1002/cbic.202300600 (2024).

31. Römer, L. & Scheibel, T. The elaborate structure of spider silk: structure and function of a natural high performance fiber. Prion 2, 154–161 10.4161/pri.2.4.7490 (2008).

32. Fan, R., et al. Sustainable Spinning of Artificial Spider Silk Fibers with Excellent Toughness and Inherent Potential for Functionalization. Advanced Functional Materials 35, 2410415 10.1002/adfm.202410415 (2025).

33. Sozcu, S., et al. Effect of Drying Methods on the Thermal and Mechanical Behavior of Bacterial Cellulose Aerogel. Gels 10, 10.3390/gels10070474 (2024).

34. Pogorelova, N., Rogachev, E., Akimbekov, N. & Digel, I. Effect of dehydration method on the micro- and nanomorphological properties of bacterial cellulose produced by Medusomyces gisevii on different substrates. Journal of Materials Science 59, 6614–6626 10.1007/s10853-024-09596-3 (2024).

35. Fuller, M. E., Andaya, C. & McClay, K. Evaluation of ATR-FTIR for analysis of bacterial cellulose impurities. Journal of Microbiological Methods 144, 145–151 10.1016/j.mimet.2017.10.017 (2018).

36. Chen, X., Knight, D. P., Shao, Z. & Vollrath, F. Conformation Transition in Silk Protein Films Monitored by Time-Resolved Fourier Transform Infrared Spectroscopy: Effect of Potassium Ions on Nephila Spidroin Films. Biochemistry 41, 14944–14950 10.1021/bi026550m (2002).

37. Zhang, H., et al. Preparation and characterization of silk fibroin as a biomaterial with potential for drug delivery. J Transl Med 10, 117 10.1186/1479-5876-10-117 (2012).

38. Andritsou, V., et al. Synthesis and Characterization of Bacterial Cellulose from Citrus-Based Sustainable Resources. ACS Omega 3, 10365–10373 10.1021/acsomega.8b01315 (2018).

39. Wei, K., Kim, B.-S. & Kim, I.-S. Fabrication and Biocompatibility of Electrospun Silk Biocomposites (2011).

40. Hubbe, M., Gardner, D. & Shen, W. Contact Angles and Wettability of Cellulosic Surfaces: A Review of Proposed Mechanisms and Test Strategies. BioResources 10, 10.15376/biores.10.4.Hubbe_Gardner_Shen (2015).

41. Abidi, W., Torres-Sánchez, L., Siroy, A. & Krasteva, P. V. Weaving of bacterial cellulose by the Bcs secretion systems. FEMS Microbiol Rev 46, 10.1093/femsre/fuab051 (2022).

42. Marmur, A. Wetting on Hydrophobic Rough Surfaces: To Be Heterogeneous or Not To Be? Langmuir 19, 8343–8348 10.1021/la0344682 (2003).

43. Chen, S.-Q., Lopez-Sanchez, P., Wang, D., Mikkelsen, D. & Gidley, M. J. Mechanical properties of bacterial cellulose synthesised by diverse strains of the genus Komagataeibacter. Food Hydrocolloids 81, 87–95 10.1016/j.foodhyd.2018.02.031 (2018).

44. Campano, C., et al. Bioengineering toughened, stretchable bacterial cellulose-based films inspired by spider silk. Carbohydrate Polymers 388, 125543 10.1016/j.carbpol.2026.125543 (2026).

45. Cannazza, P., et al. Characterization, genome analysis and genetic tractability studies of a new nanocellulose producing Komagataeibacter intermedius isolate. Scientific Reports 12, 20520 10.1038/s41598-022-24735-z (2022).

46. Florea, M., et al. Engineering control of bacterial cellulose production using a genetic toolkit and a new cellulose-producing strain. Proc Natl Acad Sci U S A 113, E3431–3440 10.1073/pnas.1522985113 (2016).

47. Schindelin, J., et al. Fiji: an open-source platform for biological-image analysis. Nat Methods 9, 676–682 10.1038/nmeth.2019 (2012).

48. Tukey, J. W. Comparing Individual Means in the Analysis of Variance. Biometrics 5, 99–114 10.2307/3001913 (1949).

