## Supplementary Information for the manuscript for "Engineering the bacterial cellulose-forming surface as a programmable protein recruitment interface"

**Supplementary Information for “Engineering the bacterial cellulose-forming surface as a programmable protein recruitment interface”**

Shuang Zhang^1^, Chengji Yang^2^, Ruxia Fan^1^, Sandra Kaabel ^2^, A. Sesilja Aranko ^1^, Markus B. Linder^1^, Rahul Mangayil ^1*^

^1^ Department of Bioproducts and Biosystems, School of Chemical Engineering, Aalto University, P.O. Box 16100, FI-02150 Espoo, Finland

^2^ Department of Chemistry and Material Sciences, School of Chemical Engineering, Aalto University, P.O. Box 16100, FI-02150 Espoo, Finland

* Corresponding author

**Contents**

**Supplementary Methods**

**Table S1** Bacterial strains, protein treatments and BC material groups.

**Table S2** Primers used in this study.

**Data S1.** Nucleotide sequences of display-protein/peptide fusions used in this study.

**Fig. S1** Agarose gel electrophoresis verification of Lpp’OmpA-based surface-display constructs.

**Fig. S2** Time-course fluorescence microscopy of Lpp’OmpA–mLime-expressing *K. intermedius*.

**Fig. S3** SpyTag-mediated recruitment of SpyCatcher–mScarlet on the *K. intermedius* cell surface.

**Fig. S4** Macroscopic imaging of bacterial cellulose pellicle formation over time.

**Fig. S5** Full SDS–PAGE gel image for analysis of pellicle-associated proteins.

**Fig. S6** Time-course fluorescence imaging of rADF3-Sp-associated bacterial cellulose pellicles.

**Fig. S7** Additional SEM images of vacuum-dried bacterial cellulose pellicles.

**Fig. S8** Additional SEM images of air-dried bacterial cellulose pellicles.

**Fig. S9** Additional SEM images of freeze-dried bacterial cellulose pellicles.

**Fig. S10** Additional SEM images of liquid-nitrogen-fractured bacterial cellulose pellicles.

**Fig. S11** Polarized optical microscopy of rADF3-Sp-associated bacterial cellulose pellicles.

**Fig. S12** Thermogravimetric analysis of rADF3-Sp-associated bacterial cellulose pellicles.

**Fig. S13** Statistical analysis of water contact angle measurements.

**Fig. S14 Bacterial cellulose pellicle formation by engineered *K. intermedius* in the presence of *E. coli* lysate.**

**Supplementary Reference**

**Supplementary Methods**

**Genetic parts, plasmids and primers**

All primers and nucleotide sequences used in this study are listed in Tables S2 and Data S1. Surface-display constructs were designed to present modular cargo domains on the surface of *K. intermedius* through fusion to the Lpp’OmpA anchor. The cargo domains included mLime, SpyTag and SilkTag. Inducible constructs were placed under the control of the P_lux_ promoter.

Plasmids were assembled using a Golden Gate cloning strategy^1^. Assembled plasmids were transformed into *Escherichia coli* TOP10 for propagation. Transformants were screened by colony PCR using construct-specific primers, and positive clones were verified by whole-plasmid sequencing. Verified plasmids were introduced into *K. intermedius* by electroporation. Positive *Komagataeibacter* transformants were selected on HS agar plates containing the appropriate antibiotics and confirmed by colony PCR.

**Preparation of purified SpyCatcher-mScarlet and silk fusion proteins**

SpyCatcher-mScarlet and SilkCatcher-rADF3-Sp-SpyTag fusion proteins were expressed in *E. coli* TOP10 and purified following a previously reported protocol. Purified proteins were analysed by SDS-PAGE, buffer-exchanged into PBS or the indicated storage buffer and quantified using before use^1^.

For SpyTag-binding assays, SpyCatcher-mScarlet was used from a 340 µM stock solution. For the preparation of rADF3-Sp-associated bacterial cellulose (BC) pellicles, purified SilkCatcher-rADF3-Sp-SpyTag was used at 5.9 g L⁻¹.

**Preparation of bacterial cellulose-derived inoculum**

BC pellicles used for inoculum preparation were produced by static cultivation in HS medium. After cultivation, pellicles were collected under sterile conditions and treated with cellulase to digest the cellulose matrix and release the cellulose-embedded cells. Cellulase digestion was performed using 2% at 30°C for 6h in HS medium. Digested samples were washed three times with sterile 1× PBS by centrifugation at 3900 rpm for 12 min. After the final wash, the pellet was collected and weighed in the wet state.

For each treatment, 0.1 g wet sample was resuspended in 400 µL PBS. rADF3-Sp-treated samples received 800 µL purified SilkCatcher-rADF3-Sp-SpyTag solution at 5.9 g L⁻¹, whereas control samples received 800 µL PBS. This corresponded to 4.72 mg protein per 0.1 g wet sample in the rADF3-Sp-treated groups. Samples were incubated at room temperature for 6 h before being used for static bacterial cellulose pellicle production.

**Fluorescence microscopy and image processing**

Fluorescence microscopy was performed using a Zeiss Axio Vert A1 (Carl Zeiss Microscopy GmbH, Jena, Germany) inverted microscope equipped with fluorescence channels suitable for mLime and mScarlet detection. For time-course imaging, pellicles were collected at 12, 24 and 36 h after inoculation. Samples compared within the same experiment were imaged using identical acquisition settings, including exposure time and illumination intensity.

Images were processed using Fiji/ImageJ^2^. Brightness and contrast were adjusted identically for images compared to the same experiment. Representative images were selected from at least three independently cultivated samples per group.

**Scanning electron microscopy sample preparation**

BC pellicles were prepared for scanning electron microscopy (SEM) after air drying, vacuum drying or freeze-drying. Air-dried samples were dried at room temperature. Vacuum-dried samples were dried at 50 °C. Freeze-dried samples were pre-frozen by liquid nitrogen and lyophilized using Labconco Freezone 2.5 Freeze Dryer ((Labconco Corporation, Kansas City, MO, USA). for one week. For fractured-interface imaging, samples were immersed in liquid nitrogen and fractured before mounting.

Samples were mounted on SEM stubs using conductive carbon tape and sputter-coated with 5 nm Au/Pd using Quorum Technologies Q 150 R Sputter Coater ((Quorum Technologies Ltd., Laughton, East Sussex, UK). SEM imaging was performed using a JEOL JSM-IT800HL Schottky field-emission scanning electron microscope (JEOL Ltd., Tokyo, Japan). at an accelerating voltage of 2 kV. Multiple regions were imaged from each independent sample to assess the reproducibility of the observed morphology.

**Polarized optical microscopy**

Polarized optical microscopy was performed using Olympus BX53M (Olympus Corporation, Tokyo, Japan) under crossed polarizers. Washed pellicles were placed on glass slides and imaged without staining. Imaging settings were kept constant across WT-BC, WT-BC/rADF3-Sp, Eng-BC and Eng-BC/rADF3-Sp samples. Representative images were obtained from at least three independent pellicles per group.

**Fourier-transform infrared spectroscopy**

Fourier-transform infrared spectroscopy was performed on freeze-dried bacterial cellulose pellicles using PerkinElmer FTIR (PerkinElmer, Inc., Waltham, MA, USA) with ATR in attenuated total reflectance mode. Spectra were collected over 4000-500 cm^-1^ and processed by baseline correction and normalization. The amide I/II region was examined to assess rADF3-Sp-associated protein features.

**X-ray diffraction**

X-ray diffraction measurements were performed on dried pellicles using Bruker Advance D8 Powder XRD (Bruker AXS GmbH, Karlsruhe, Germany) with Cu Kα radiation. Diffraction patterns were collected over a 2θ range of 10°to 40° with a step size of with and a scan rate of 1s. Crystallinity index values were calculated using Crystallinity index (CrI) = 100 × (*I*_200_ − *I*_AM_)/*I*_200_^3^_._ Characteristic BC diffraction peaks were compared among WT-BC, WT-BC/rADF3-Sp, Eng-BC and Eng-BC/rADF3-Sp samples.

**SDS-PAGE analysis of pellicle-associated proteins**

Washed BC pellicles were digested with cellulase at 30 °C for 6 h to release pellicle-associated proteins. The soluble fractions were collected, mixed with SDS-PAGE loading buffer, heated at 95 °C for 5 min and analysed by SDS-PAGE followed by Coomassie Brilliant Blue staining. Purified SilkCatcher-rADF3-Sp-SpyTag was loaded as a reference protein. WT-BC, WT-BC/rADF3-Sp, Eng-BC and Eng-BC/rADF3-Sp samples were compared to assess rADF3-Sp-associated protein retention after washing.

**Thermogravimetric analysis**

Thermogravimetric analysis was performed using TA Instruments TGA 5500 (TA Instruments, New Castle, DE, USA). Dried pellicle samples were heated from30° to 800°at a heating rate of 10°/min under nitrogen. Residual mass was calculated from the remaining mass at the final temperature. TGA profiles were compared among rADF3-Sp-associated pellicles and their corresponding controls.

**Unconfined compression testing and analysis**

Hydrated BC pellicles were tested using an Instron 4204 universal (Instron Corporation, Norwood, MA, USA) testing machine. Samples were maintained in the hydrated state before measurement. The sample height was approximately 3 mm, and the exact dimensions of each sample were recorded before testing. A preload of 0.2 N was applied before compression. Samples were compressed at 0.02 mm s⁻¹ to 60% strain unless otherwise stated.

Raw force–displacement data were exported for analysis. Engineering strain was calculated as ε = Δh/h₀, where Δh is the compressive displacement and h₀ is the initial hydrated sample height. Engineering stress was calculated as σ = F/A₀, where F is the measured compressive force and A₀ is the initial cross-sectional area of the pellicle.^4, 5, 6^ Stress–strain curves were generated for each replicate, and compressive stresses at 30%, 50% and 60% strain were extracted from the corresponding curves. The apparent compressive modulus was calculated from the slope of the stress–strain curve between 0 and 10% strain. Data are reported as mean ± standard deviation (s.d.), with the number of biological or technical replicates indicated in the figure captions.

**Water contact angle measurements**

Static water contact angles were measured using a Theta Flex optical tensiometer (Attension, Biolin Scientific AB, Gothenburg, Sweden). Dried pellicles were placed on the sample stage, and 5µL deionized water droplets were deposited onto the sample surface. Images were recorded immediately after droplet deposition and, where applicable, over 10s to monitor droplet spreading. Contact angles were calculated using OneAttension (Attension, Biolin Scientific AB, Gothenburg, Sweden).

Measurements were performed at multiple positions on each sample. Three independent pellicles were measured per group. Because BC samples can absorb water rapidly, the time point used for contact angle comparison was kept constant across all groups.

**Statistical analysis and reproducibility**

Statistical analyses were performed using GraphPad Prism version 10 and Origin 2021. Data are presented as mean ± s.d. unless otherwise stated. Comparisons among multiple groups were performed using one-way ANOVA followed by Tukey’s multiple-comparison test^7^. Biological replicates refer to independently prepared bacterial cultures or independently produced pellicles, whereas technical replicates refer to repeated measurements from the same biological sample. Replicate numbers are indicated in the corresponding figure legends. Representative microscopy images were selected from at least three independent samples or experiments.

**BC pellicle formation in lysate-containing medium**

To assess whether *E. coli* lysate affected BC production, *K. intermedius* cells were inoculated into HS medium containing different volume fractions of *E. coli* lysate. Wild-type (WT) *K. intermedius* cultured in standard HS medium was used as reference control. Engineered *K. intermedius* cells were cultured in HS medium without lysate or supplemented with 5%, 10%, 15%, 20%, 30% or 50% *E. coli* lysate. The initial OD₇₀₀_nm_ was adjusted to 0.1 for all groups. Cultures were incubated statically at 30 °C for 5 days, after which pellicle formation was photographed.

**Table S1 Bacterial strains, protein treatments and BC material groups.**

| Strain name | Surface module | Protein treatment | Meaning |
| --- | --- | --- | --- |
| WT-BC | None | PBS | BC pellicle produced by WT cells without rADF3-Sp exposure |
| WT-BC/rADF3-Sp | None | SilkCatcher-rADF3-Sp-SpyTag | BC pellicle produced by WT cells exposed to the silk-derived fusion protein without a defined surface docking site |
| Eng-BC | Lpp’OmpA-SilkTag | PBS | BC pellicle produced by SilkTag-displaying engineered cells without rADF3-Sp exposure |
| Eng-BC/rADF3-Sp | Lpp’OmpA-SilkTag | SilkCatcher-rADF3-Sp-SpyTag | BC pellicle produced after SilkTag-mediated recruitment of the silk-derived fusion protein before cellulose biosynthesis |

“Eng” denotes *K. intermedius* expressing Lpp’OmpA-SilkTag; “BC” denotes the resulting bacterial cellulose pellicle; “/rADF3-Sp” denotes pre-incubation with SilkCatcher-rADF3-Sp-SpyTag before static cellulose biosynthesis

**Table S2** **Primers used in this study.**

| Primer | Sequence (5'→3') |
| --- | --- |
| *Primers used to verify Lpp’OmpA-mLime construct* | |
| EY-F-mlime | tatagtcctgtcgggtttcgcc |
| EY-R-mlime | ccggtgagcgtgggtcccgcggtatc |
| L1-F-mlime | agggcggcggatttgtcc |
| L1-R-mlime | gttctgaggtcattactgg |
| L2-F-mlime | agggcggcggatttgtcc |
| L2-R-mlime | gttctgaggtcattactgg |
| *Primers used to construct and verify the Lpp’OmpA-SpyTag construct* | |
| Spy-F | gttatggttgatgcatacaaaccgaccaaaTAACATCATCACCACCATCATttctttatt |
| Spy-R | tatgcatcaaccataacaatatgtgcTGAACCGCCACCACCGCTACCACCACCACCTTGA |
| Test-Spy-F | AATCATGATACCGGTGTTAGTC |
| Test-Spy-R | gaaaccttaacgctatgg |
| *Primers used to construct and verify the Lpp’OmpA-SilkTag construct* | |
| Silk-F | TTGCGTTCCAGGTGAGCCAGGACGACGTTAAACAGCCGGTGGTACCGACCTAACATCATCACCACCATCATttctttatt |
| Silk-R | CTGGCTCACCTGGAACGCAACTTCCGGTTTGATACCTGAACCGCCACCACCGCTACCACCACCACCTTGA |
| Test-Silk-F | AATCATGATACCGGTGTTAGTC |
| Test-Silk-R | gaaaccttaacgctatgg |

**Data S1**

Lpp’OmpA sequence: ATGAAAGCCACCAAACTGGTTCTGGGTGCAGTTATTCTGGGTAGCACCCTGCTGGCAGGTTGTAGCAGCAATGCAAAAATTGATCAGGGCATTAATCCGTATGTGGGTTTTGAAATGGGTTATGATTGGCTGGGTCGTATGCCGTATAAAGGTAGCGTTGAAAATGGTGCATATAAAGCACAGGGTGTTCAGCTGACCGCAAAACTGGGTTATCCGATTACCGATGATCTGGATATCTATACCCGTTTAGGTGGTATGGTTTGGCGTGCAGATACCAAAAGCAATGTGTATGGCAAAAATCATGATACCGGTGTTAGTCCGGTTTTTGCCGGTGGTGTTGAATATGCAATTACACCGGAAATTGCAACCCGTCTGGAATATCAGTGGACCAATAACATTGGTGATGCACATACCATTGGCACCCGTCCGGATAATGGTGTGGAAAATCTGTATTTTCAAGGT

SpyTag sequence:

GCACATATTGTTATGGTTGATGCATACAAACCGACCAAA

SpyCatcher sequence:

ATGGGTGCAATGGTTACCACACTGAGCGGTCTGAGTGGTGAACAGGGTCCGAGCGGTGATATGACCACCGAAGAAGATAGCGCAACCCATATCAAATTTAGCAAACGTGATGAAGATGGTCGTGAACTGGCAGGCGCAACAATGGAACTGCGTGATAGCAGCGGTAAAACCATTAGCACCTGGATTAGTGATGGTCACGTGAAAGATTTTTATCTGTATCCGGGTAAATATACCTTCGTTGAAACCGCAGCACCGGATGGTTATGAAGTTGCAACCGCAATTACCTTTACCGTGAATGAACAAGGTCAGGTTACCGTTAATGGTGAAGCAACCAAAGGTGATGCACATACCCTCGAGTGA

SilkTag sequence:

GGTATCAAACCGGAAGTTGCGTTCCAGGTGAGCCAGGACGACGTTAAACAGCCGGTGGTACCGACCTAA

SilkCatcher sequence:

ACCTATACTATCGAACTGACCAAACACGATGCCGCGACCAAAGCGGTTCTGGCTGGTGCTGTTTATGAGCTGCAGGACTCCACCGGTAAAGTGATCCAGACCGGTCTGACCACCGACTCTCAGGGCCAGCTGATCGTTAAGAACCTGCGTGCGGGCGACTATCAGTTCGTGGAAACCAAAGCACCGCTGGGCTATGAGCTGAACACCACCCCGGTAAAATTCACCCTGGGT

rADF3-Sp sequence:

GGTCCTGGTCAGCAGGGTCCGGGTCAACAAGGACCTGGACAGCAAGGACCGTATGGTCCAGGTGCATCAGCTGCAGCCGCAGCAGCGGGTGGTTATGGTCCGGGAAGCGGTCAGCAAGGCCCTTCACAACAGGGACCAGGCCAACAGGGTCCTGGCGGTCAAGGTCCTTATGGACCTGGTGCTTCTGCTGCGGCAGCGGCTGCCGGTGGCTATGGCCCTGGTAGTGGCCAGCAAGGGCCTGGTGGCCAGGGTCCATATGGCCCAGGTTCTAGTGCCGCAGCTGCTGCTGCAGGCGGTAATGGACCGGGTTCAGGACAACAAGGTGCAGGGCAGCAAGGTCCCGGACAACAGGGTCCAGGTGGTAGTGCAGCAGCGGCAGCAGCTGGCGGATATGGACCAGGTAGTGGGCAACAAGGCCCAGGTCAACAAGGGCCAGGGGGTCAAGGCCCATACGGTCCGGGTGCTTCCGCAGCCGCAGCTGCAGCAGGCGGTTACGGTCCTGGTAGTGGTCAAGGTCCAGGCCAGCAAGGACCAGGTGGACAAGGGCCTTACGGACCAGGCGCATCTGCGGCAGCAGCAGCCGCAGGGGGATATGGTCCTGGTTCAGGGCAGCAGGGACCAGGTCAGCAAGGTCCAGGTCAGCAGGGACCTGGGGGTCAGGGACCTTACGGTCCTGGCGCAAGTGCAGCTGCAGCGGCAGCGGGTGGCTACGGACCGGGTTATGGCCAGCAGGGACCGGGACAGCAGGGACCCGGTGGACAGGGTCCGTATGGACCGGGTGCAAGTGCAGCATCAGCAGCAAGTGGTGGTTACGGACCTGGCTCAGGACAGCAAGGCCCTGGCCAACAAGGCCCTGGCGGACAGGGACCCTATGGGCCAGGTGCCAGCGCTGCAGCAGCCGCAGCCGGTGGATACGGTCCAGGCTCTGGTCAACAAGGTCCTGGGCAACAAGGTCCTGGCCAGCAGGGTCCAGGACAGCAAGGGCCTGGCGGTCAAGGACCGTACGGACCGGGTGCCAGCGCAGCGGCTGCAGCGGCAGGCGGTTATGGTCCAGGATCAGGCCAGCAAGGTCCGGGTCAGCAAGGCCCAGGGCAGCAAGGACCGGGTCAACAGGGACCGGGTCAGCAGGGTCCTGGGCAACAGGGTCCGGGACAACAGGGACCAGGTCAACAAGGACCGGGTCAACAAGGTCCAGGTGGTCAGGGTGCATATGGTCCTGGCGCTTCAGCAGCAGCAGGGGCTGCAGGGGGTTATGGCCCAGGTAGCGGTCAGCAGGGACCCGGACAACAAGGCCCTGGACAACAGGGTCCCGGTCAGCAAGGGCCAGGCCAACAAGGTCCAGGACAACAAGGACCAGGGCAGCAGGGTCCAGGCCAACAAGGCCCTTATGGTCCGGGTGCCAGTGCTGCGGCAGCGGCAGCTGGGGGTTATGGTCCAGGCTCTGGACAGCAGGGACCTGGCCAGCAAGGACCTGGGCAGCAAGGGCCAGGCGGTCAG

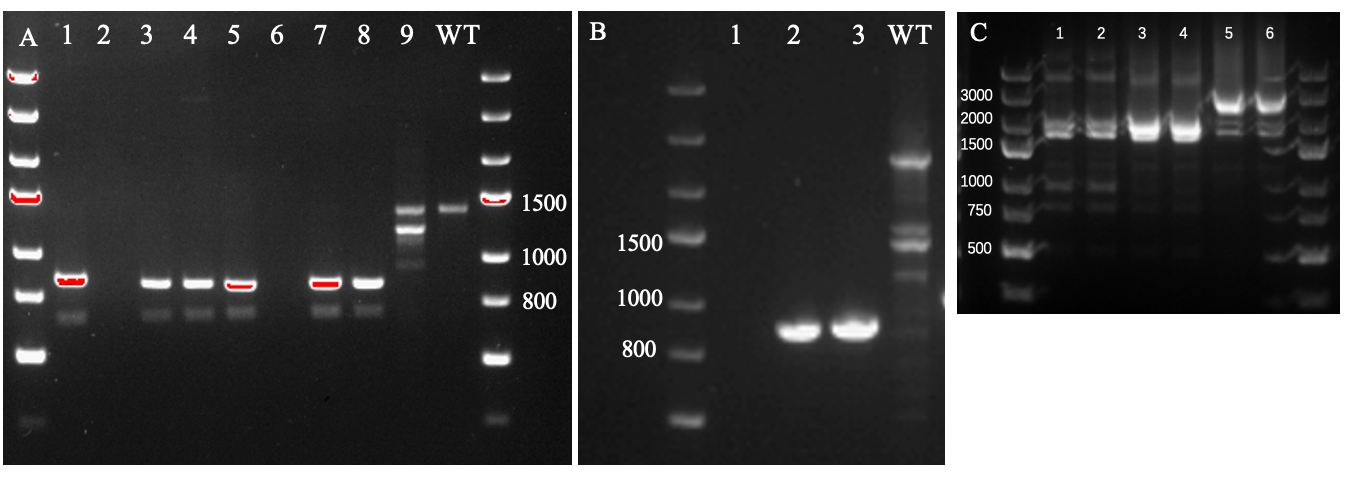

**Fig. S1 Agarose gel electrophoresis verification of Lpp’OmpA-based surface-display constructs**

A,B, Colony PCR verification of *K. intermedius* transformants carrying the P_lux_-controlled Lpp’OmpA-SpyTag construct. Numbered lanes correspond to independent transformant colonies, and WT denotes the wild-type control. The expected amplicon size was 837 bp.
C, PCR verification of Lpp’OmpA-based surface-display constructs. Lanes 1-2, Lpp’OmpA-SilkTag, 1988 bp; lanes 3-4, Lpp’OmpA-SpyTag, 2261 bp; lanes 5-6, Lpp’OmpA-mLime, 2633 bp. DNA ladders are shown on both sides.

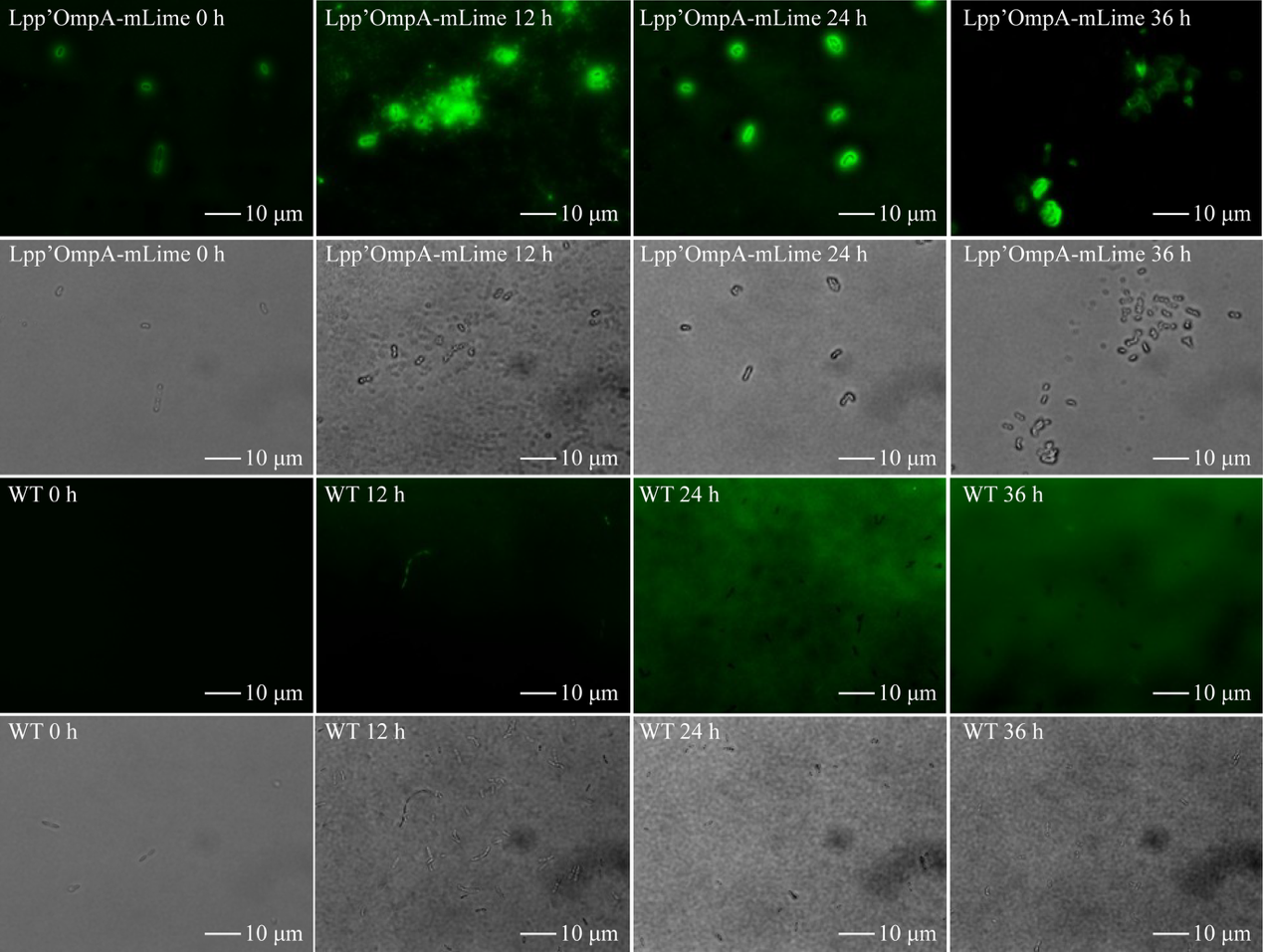

**Fig. S2 Time-course fluorescence microscopy of Lpp’OmpA-mLime-expressing *K. intermedius*.**

Lpp’OmpA-mLime-expressing cells showed green fluorescence with a predominantly peripheral pattern throughout the time course, consistent with cell-envelope-associated localization, whereas WT cells showed only background signal under the same imaging conditions. Scale bars, 10 µm.

Circular features visible in the transmitted-light images are optical artifacts from the microscope setup and do not represent sample structures.

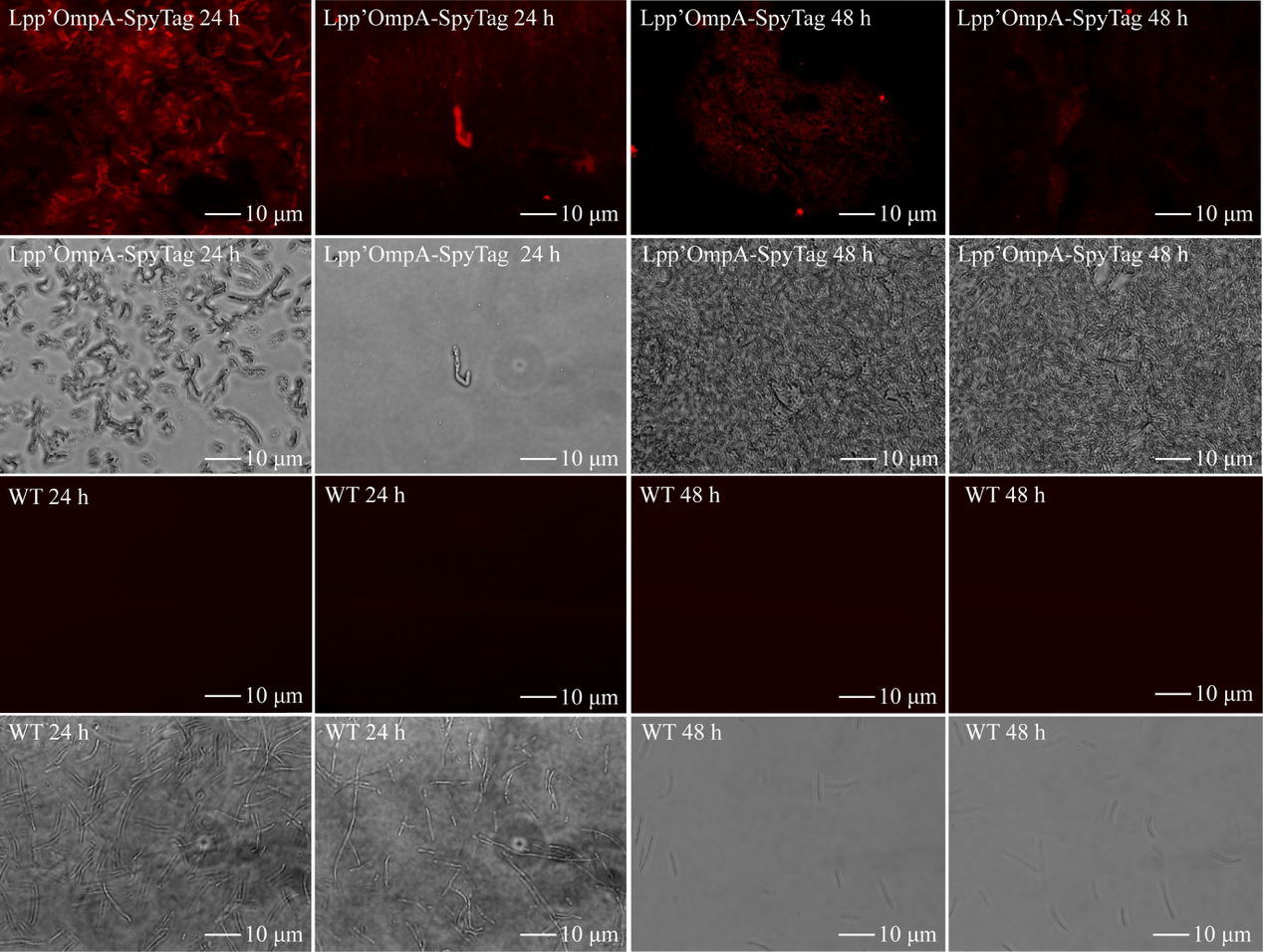

**Fig. S3 Fluorescence microscopy validation of SpyTag-mediated protein recruitment in *K. intermedius*.**

Representative fluorescence and corresponding bright-field images of Lpp’OmpA-SpyTag-expressing *K. intermedius* cells and WT controls after incubation with SpyCatcher-mScarlet. Images were collected from 24 h and 48 h cultures. Lpp’OmpA-SpyTag-expressing cells showed red fluorescence after SpyCatcher-mScarlet treatment, whereas WT cells showed only background signal under the same imaging conditions. Scale bars, 10 µm.

Circular features visible in the transmitted-light images are optical artifacts from the microscope setup and do not represent sample structures.

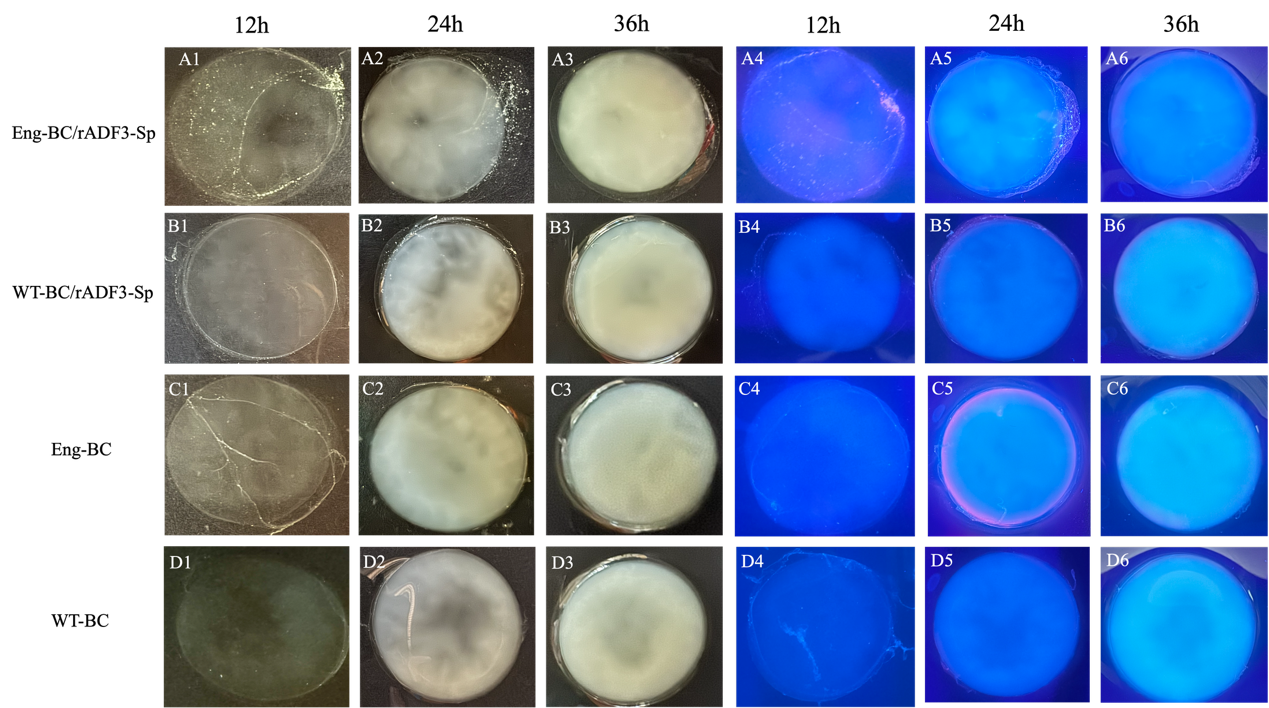

**Fig. S4 Macroscopic imaging of BC pellicle formation over time.**

Representative bright-field and fluorescence images of BC pellicles formed by the four experimental groups during static cultivation. Rows indicate sample groups: A, Eng-BC/rADF3-Sp; B, Eng-BC; C, WT-BC/rADF3-Sp; D, WT-BC. Columns 1-3 show bright-field images collected at 12, 24 and 36 h, respectively; columns 4-6 show the corresponding fluorescence images collected at the same time points. Eng-BC/rADF3-Sp showed more pronounced fluorescence-associated signal during pellicle formation compared with the control groups.

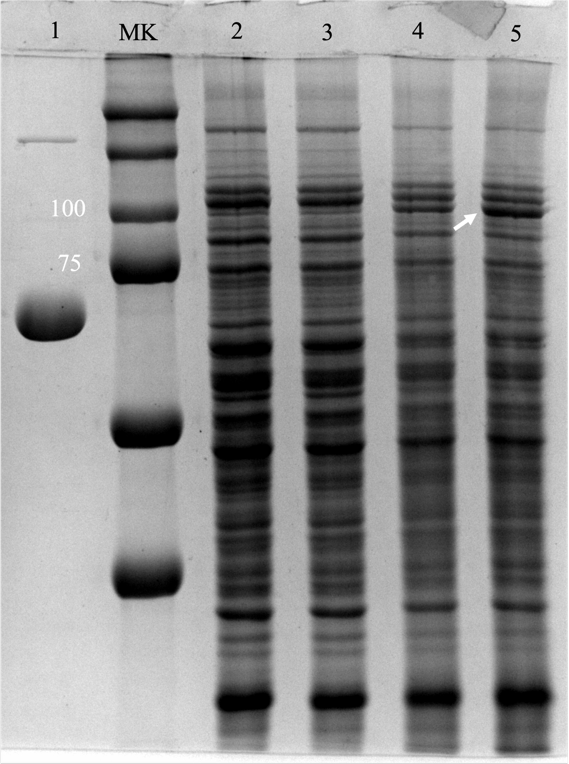

**Fig. S5 Full SDS–PAGE gel image for analysis of pellicle-associated proteins.**

Uncropped SDS–PAGE gel image corresponding to the cropped gel shown in Fig. 2C. Washed bacterial cellulose pellicles were digested with cellulase to release pellicle-associated proteins, and extracts from WT-BC, WT-BC/rADF3-Sp, Eng-BC and Eng-BC/rADF3-Sp samples were analysed by SDS–PAGE followed by Coomassie Brilliant Blue staining. Purified SilkCatcher–rADF3-Sp–SpyTag was loaded as a reference protein. The boxed or indicated region corresponds to the cropped area shown in the main Figure. 2

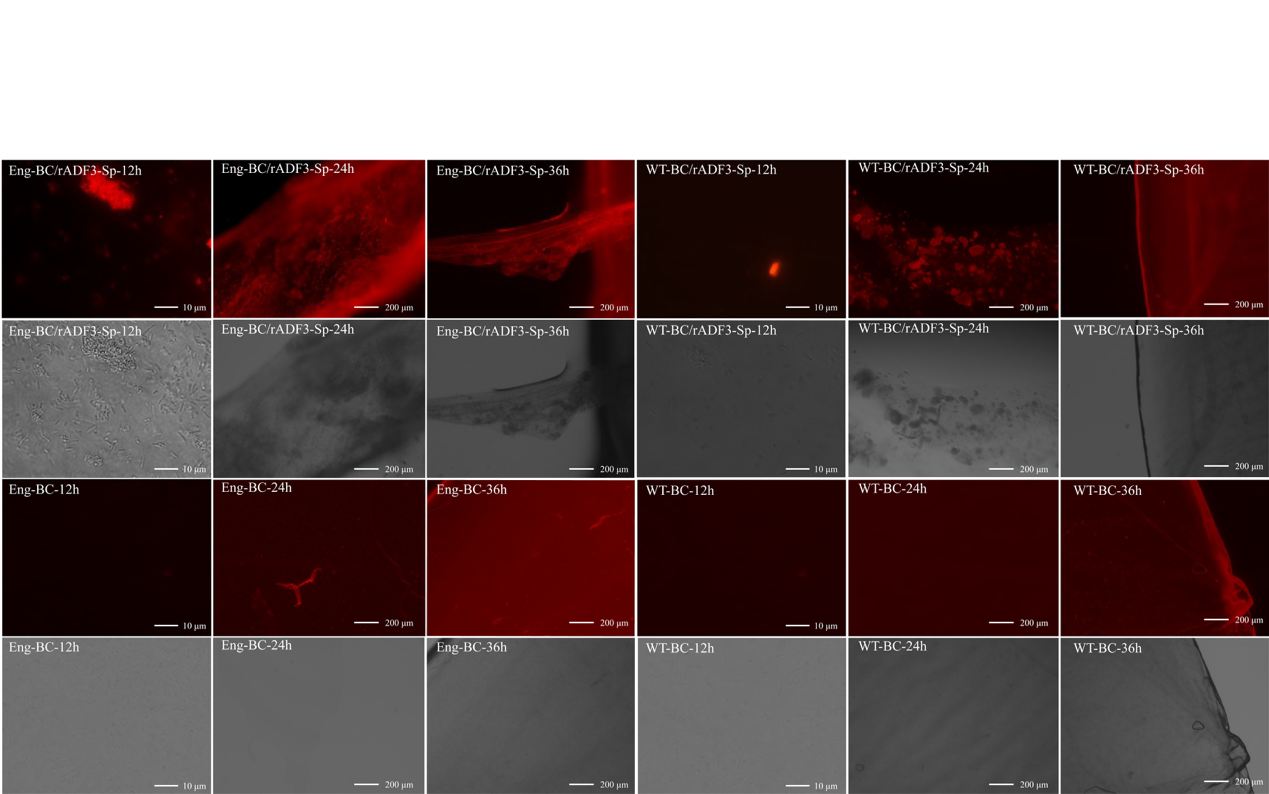

**Fig. S6 Time-course fluorescence imaging of rADF3-Sp-associated BC cellulose pellicles.**

Representative fluorescence and corresponding bright-field images of washed bacterial cellulose pellicles after SpyCatcher-mScarlet labelling. Images were collected from Eng-BC/rADF3-Sp, WT-BC/rADF3-Sp, Eng-BC and WT-BC samples at 12, 24 and 36 h after static cultivation. Eng-BC/rADF3-Sp showed more pronounced red fluorescence over the time course than the control groups, supporting retention of SpyTag-containing rADF3-Sp-associated protein in the engineered pellicles. Scale bars, 10 µm for 12 h images and 200 µm for 24 and 36 h images.

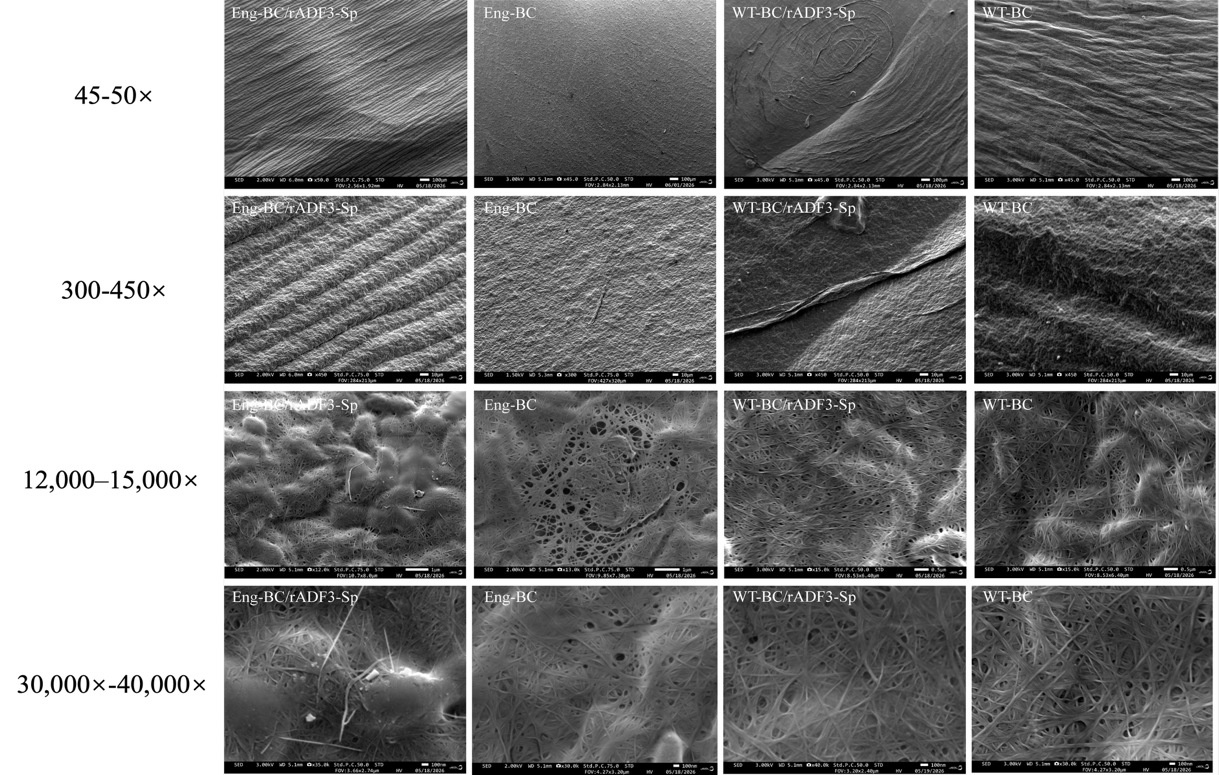

**Fig. S7 SEM images of vacuum-dried BC pellicles.**

Representative SEM images of vacuum-dried Eng-BC/rADF3-Sp, Eng-BC, WT-BC/rADF3-Sp and WT-BC pellicles acquired from multiple surface regions and magnifications. Images are arranged by approximate magnification from top to bottom 50×, 450×, 12,000–15,000× and 30,000×. These images provide additional views of the surface morphologies shown in Fig. 3A. Scale bars are indicated in the images.

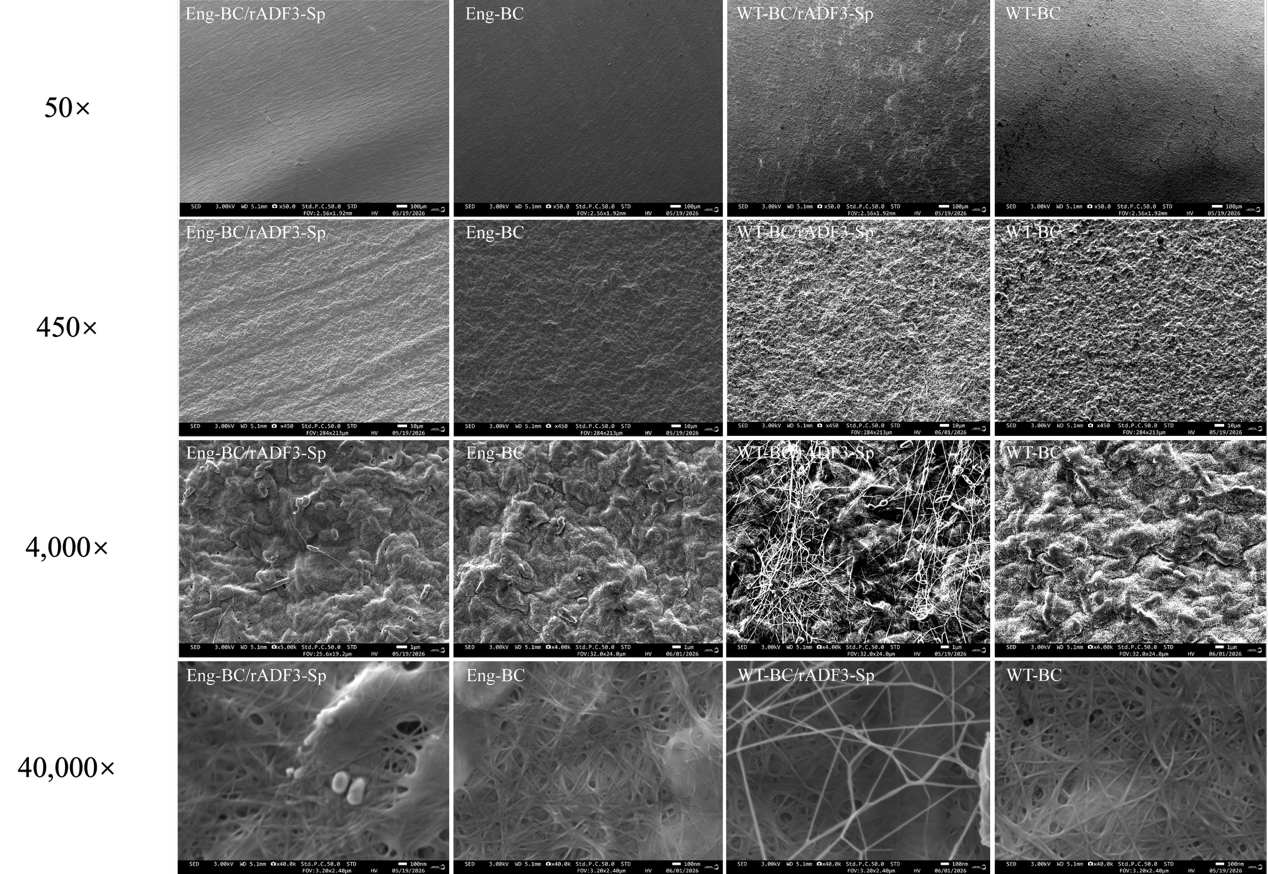

**Fig. S8 SEM images of air-dried BC pellicles**

Representative SEM images of air-dried Eng-BC/rADF3-Sp, Eng-BC, WT-BC/rADF3-Sp and WT-BC pellicles acquired from multiple surface regions and magnifications. Images are arranged by approximate magnification from top to bottom 50×, 450×, 4,000× and 40,000×. These images provide additional views of the surface morphologies of air-dried bacterial cellulose pellicles and illustrate the influence of drying method on the observed surface structure. Scale bars are indicated in the images.

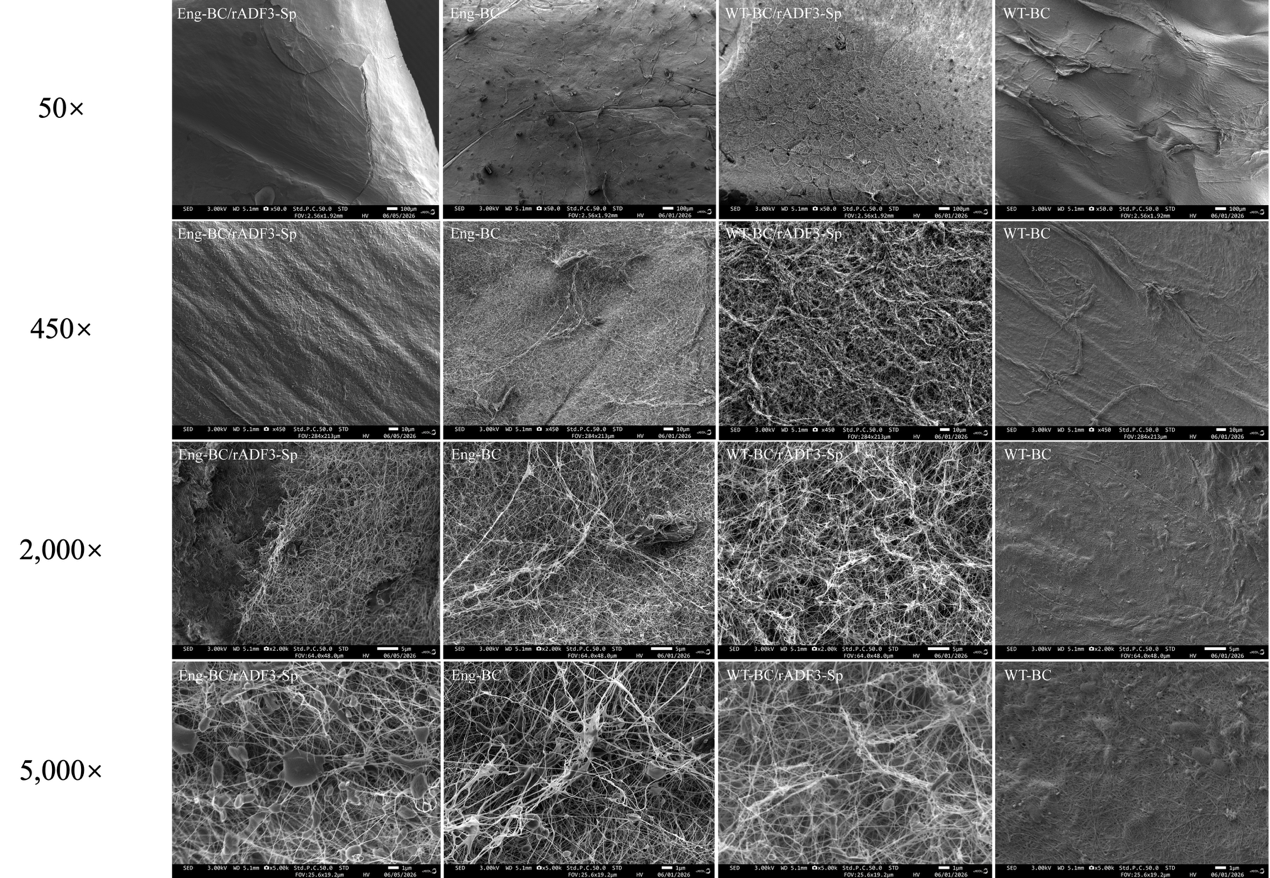

**Fig. S9 SEM images of freeze-dried BC pellicles.**

Representative SEM images of freeze-dried flat surfaces of Eng-BC/rADF3-Sp, Eng-BC, WT-BC/rADF3-Sp and WT-BC pellicles acquired from multiple surface regions and magnifications. These images provide additional views of the freeze-dried surface morphology and illustrate the influence of sample preparation on the observed bacterial cellulose network structure. Scale bars are indicated in the images.

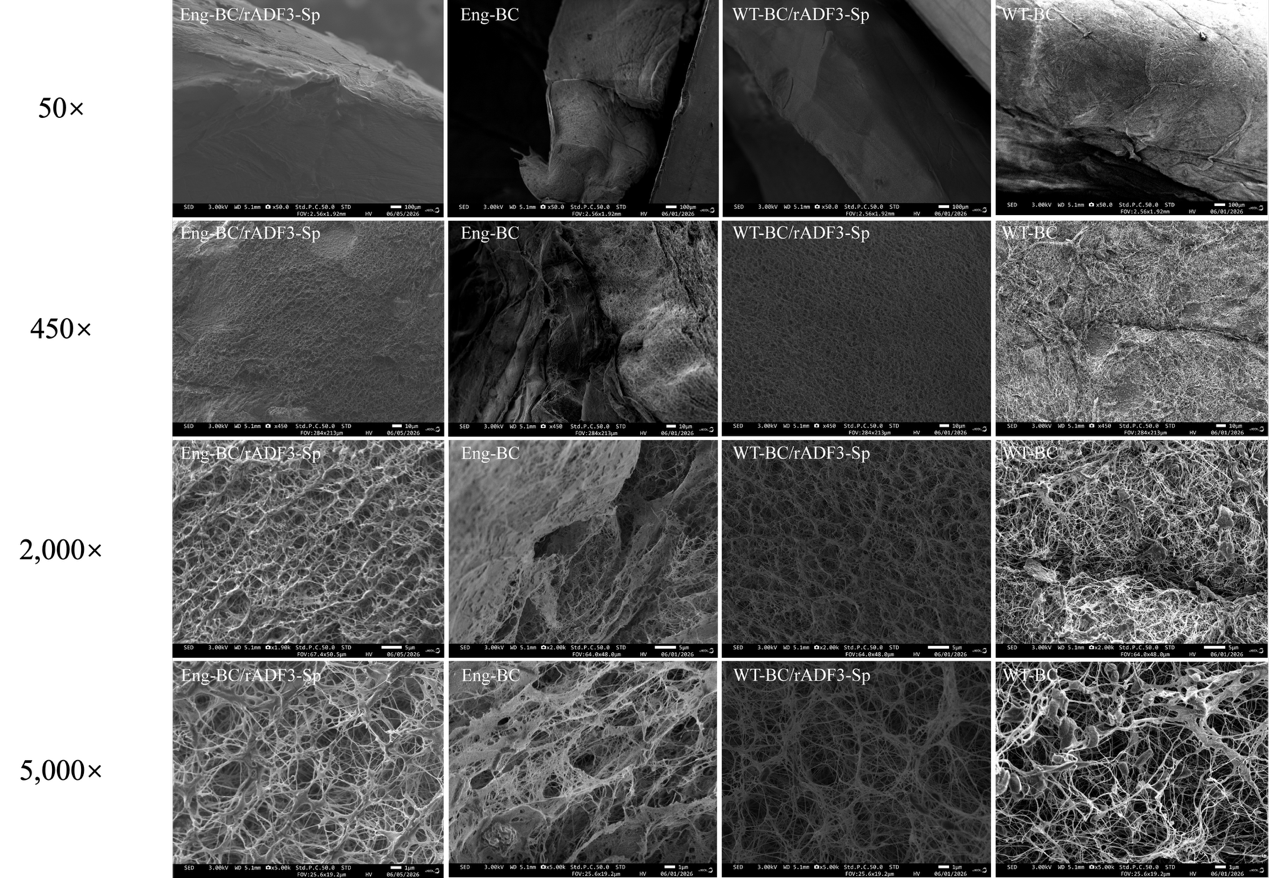

**Fig. S10 SEM images of liquid-nitrogen-fractured BC pellicles**

Representative SEM images of liquid-nitrogen-fractured Eng-BC/rADF3-Sp, Eng-BC, WT-BC/rADF3-Sp and WT-BC pellicles acquired from multiple fractured regions and magnifications. These images provide additional views of the internal fractured-interface morphology shown in Fig. 3B and support comparison of the cellulose-associated fibrous networks among the four groups. Scale bars are indicated in the images.

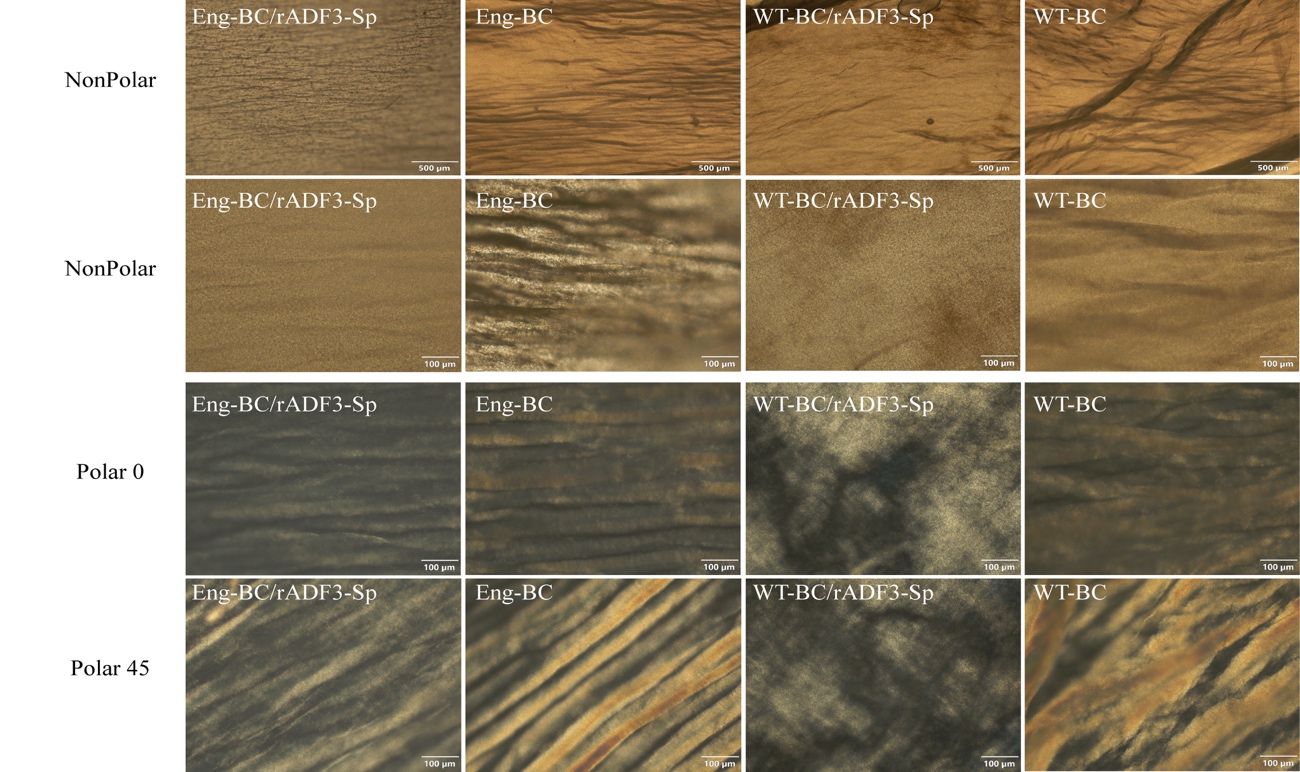

**Fig. S11 Polarized optical microscopy of BC pellicles.**

Representative optical microscopy images of Eng-BC/rADF3-Sp, Eng-BC, WT-BC/rADF3-Sp and WT-BC pellicles acquired under non-polarized and polarized imaging conditions. Non-polarized images were collected at two magnifications, and polarized images were collected at analyser angles of 0° and 45°. Eng-BC/rADF3-Sp showed continuous band-like optical features under polarized imaging, consistent with altered mesoscale organization after SilkTag-mediated rADF3-Sp recruitment. Scale bars, 500 µm for low-magnification non-polarized images and 100 µm for the remaining images.

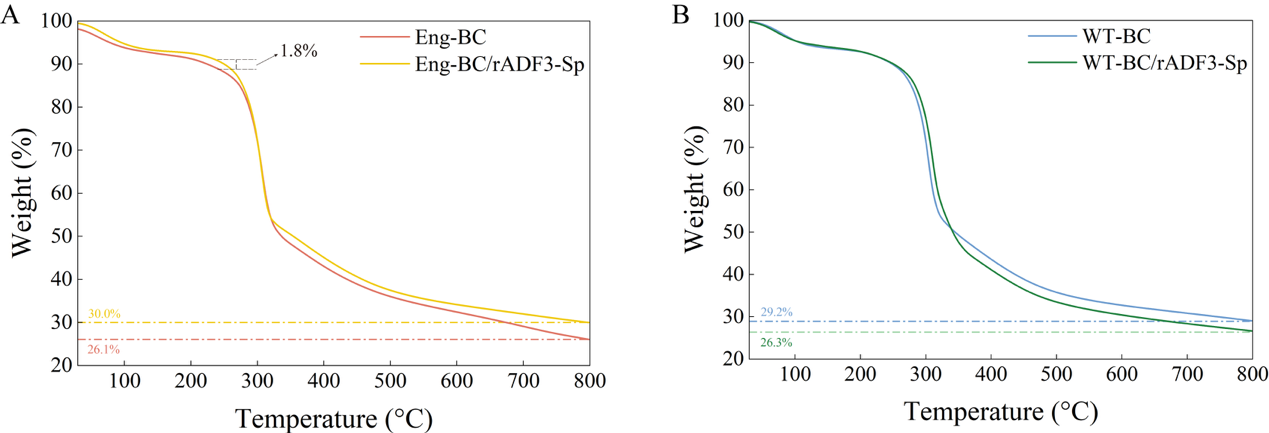

**Fig. S12 Thermogravimetric analysis of BC pellicles.**

**A**, TGA curves of Eng-BC and Eng-BC/rADF3-Sp dried pellicles. **B**, TGA curves of WT-BC and WT-BC/rADF3-Sp dried pellicles.

All samples showed a minor initial weight loss below approximately 120 °C and a major degradation step around 280–330 °C. Eng-BC/rADF3-Sp showed a slightly higher residual mass than Eng-BC at 800 °C, approximately 30.0% versus 26.1%, whereas WT-BC/rADF3-Sp showed a slightly lower residual mass than WT-BC, approximately 26.3% versus 29.2%. These residual-mass differences suggest modest changes in the thermal response of the dried pellicles after rADF3-Sp treatment or recruitment.

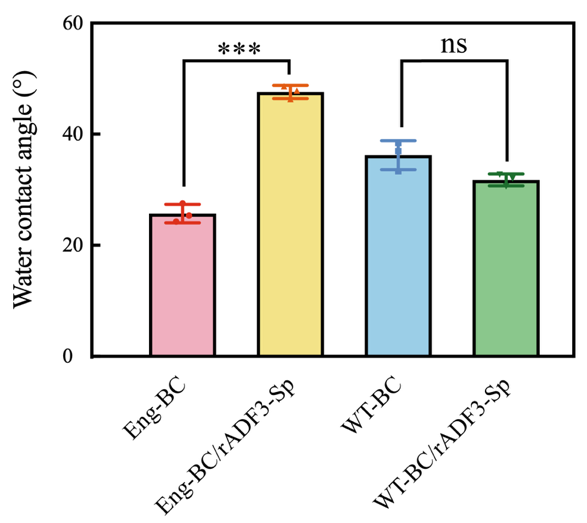

**Fig. S13 Statistical analysis of water contact angle measurements.**

Static water contact angles of dried Eng-BC, Eng-BC/rADF3-Sp, WT-BC and WT-BC/rADF3-Sp pellicles. Eng-BC/rADF3-Sp showed a significantly higher water contact angle than Eng-BC, whereas WT-BC/rADF3-Sp was not significantly different from WT-BC. Data are shown as mean ± SD. Statistical analysis was performed using one-way ANOVA followed by Tukey’s multiple-comparison test; ***P < 0.001; ns, not significant.

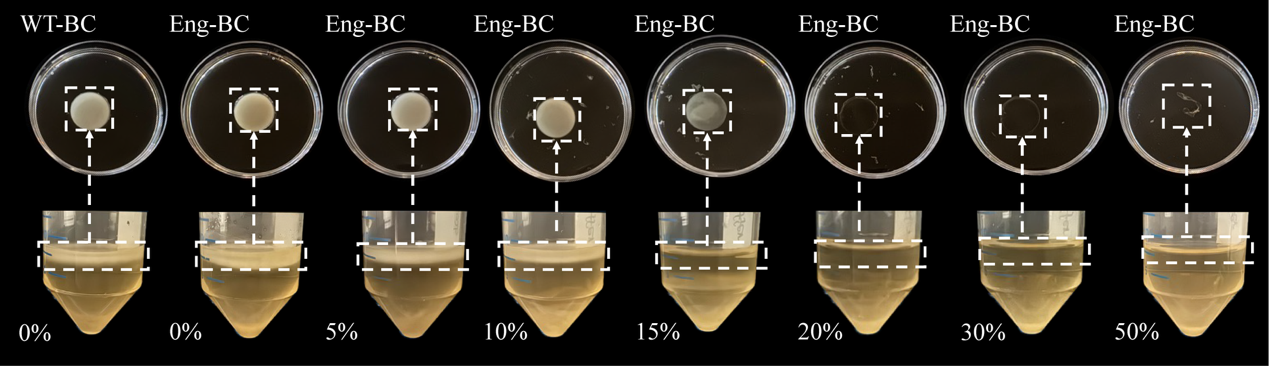

**Fig. S14 BC pellicle formation by engineered *K. intermedius* in the presence of *E. coli* lysate.**

Representative images of BC pellicles formed after 5 days of static cultivation. WT K. intermedius cultured in standard HS medium was used as reference control. Engineered K. intermedius cells were cultured in HS medium without lysate. The initial OD₇₀₀ was adjusted to 0.1 for all groups. Pellicle formation under lysate-containing conditions indicates that the lysate environment did not completely prevent BC biosynthesis.

**Supplementary References**

1.	Goosens, V. J.*, et al.* Komagataeibacter Tool Kit (KTK): A Modular Cloning System for Multigene Constructs and Programmed Protein Secretion from Cellulose Producing Bacteria. *ACS Synthetic Biology* **10**, 3422–3434 <https://doi.org/10.1021/acssynbio.1c00358> (2021).

2.	Schindelin, J.*, et al.* Fiji: an open-source platform for biological-image analysis. *Nature Methods* **9**, 676–682 <https://doi.org/10.1038/nmeth.2019> (2012).

3.	Salem, K. S.*, et al.* Comparison and assessment of methods for cellulose crystallinity determination. *Chemical Society Reviews* **52**, 6417–6446 <https://doi.org/10.1039/d2cs00569g> (2023).

4.	Oyen, M. L. Mechanical characterisation of hydrogel materials. *International Materials Reviews* **59**, 44–59 <https://doi.org/10.1179/1743280413Y.0000000022> (2014).

5.	Lopez-Sanchez, P.*, et al.* Micromechanics and Poroelasticity of Hydrated Cellulose Networks. *Biomacromolecules* **15**, 2274–2284 <https://doi.org/10.1021/bm500405h> (2014).

6.	Treesuppharat, W., Rojanapanthu, P., Siangsanoh, C., Manuspiya, H. & Ummartyotin, S. Synthesis and characterization of bacterial cellulose and gelatin-based hydrogel composites for drug-delivery systems. *Biotechnology Reports* **15**, 84–91 <https://doi.org/https://doi.org/10.1016/j.btre.2017.07.002> (2017).

7.	Tukey, J. W. Comparing Individual Means in the Analysis of Variance. *Biometrics* **5**, 99–114 <https://doi.org/10.2307/3001913> (1949).
